# A pan-respiratory virus attachment inhibitor with high potency in human airway models and in vivo

**DOI:** 10.1101/2025.01.25.634704

**Authors:** Gregory Mathez, Paulo Jacob Silva, Vincent Carlen, Charlotte Deloizy, Clémentine Prompt, Anaïs Hubart, Joana Rocha-Pereira, Marie Galloux, Ronan Le Goffic, Francesco Stellacci, Valeria Cagno

## Abstract

Respiratory viruses can cause severe infections, including bronchiolitis and pneumonia, often leading to hospitalization or death. Due to their ease of transmission, they are also scrutinized for their pandemic potential. No broad-spectrum antiviral is currently available. However most respiratory viruses use sialic acid or heparan sulfates as attachment receptors. Here, we report the identification of a pan-respiratory antiviral strategy based on mimicking both glycans. We synthesized and characterized a unique modified cyclodextrin that simultaneously mimics heparan sulfate and sialic acid. This novel compound demonstrated broad-spectrum antiviral activity against important human pathogens: parainfluenza virus 3, respiratory syncytial virus, influenza virus H1N1, SARS-CoV-2. Additionally, the compound is active against different avian strains of influenza virus, revealing its importance for pandemic preparedness. The compound retains broad-spectrum activity in ex vivo models of respiratory tissues and in vivo experiments against RSV and Influenza virus, using both prophylactic and therapeutic strategies. These findings represent a significant step forward in the development of future treatments and preventive measures for respiratory viral infections.

**Significance Statement:** Following the SARS-CoV-2 pandemic, Influenza A H5N1 has become widespread in poultry and has already begun to spill over into mammals, posing a potential risk for the next pandemic. Indeed, respiratory viruses represent a major threat for future global health crises. Unfortunately, there is a significant lack of broad-spectrum antivirals available for such scenarios. To address this gap, our study developed a single molecule with the capacity to inhibit a wide range of clinically relevant human respiratory viruses including avian strains of influenza virus. This antiviral demonstrated verified efficacy not only in cellular systems but also in human-derived respiratory tissues and animal models.

## Introduction

Respiratory infections remain one of the leading causes of global mortality, driven primarily by RNA viruses from various families, including Coronaviridae (e.g., SARS-CoV-2), Picornaviridae (e.g., Enterovirus, Rhinovirus), Orthomyxoviridae (e.g., Influenza A virus (IAV)), Paramyxoviridae (e.g., hPIV), and Pneumoviridae (e.g., RSV). These viruses are transmitted via aerosols and droplets, infecting the respiratory tract, with seasonal outbreaks influenced by social behaviors and environmental conditions [1, 2]. Vulnerable populations such as children, the elderly, and immunocompromised individuals are particularly prone to severe lower respiratory tract infections, including bronchiolitis and pneumonia.

Respiratory viruses are under close surveillance for their pandemic potential, linked to their high transmissibility and capacity for rapid mutation, enabling them to adapt to new environments. In December 2019, SARS-CoV-2 was first identified in China [3]. As the pandemic unfolded, SARS-CoV-2 evolved into a major pathogen. By late 2020, the FDA approved the first antiviral treatment, followed shortly by the rollout of vaccines—an unprecedented achievement within a year of global spread. More recently, the cross-species transmission of avian IAV H5N1 to mammals has heightened the urgency to contain this pathogen and prevent human infection [4, 5], while other avian influenza strains continue to pose a significant zoonotic threat. Despite the threat posed by respiratory viruses, approved treatments are lacking for most of them. Developing broad-spectrum antivirals is critical for future pandemic preparedness and for providing potential therapies. However, finding effective broad-spectrum antivirals relies on conserved targets present in or used by diverse viruses without toxicity for host cells.

Among the inhibitors investigated for antiviral development, those targeting viruses’ attachment to the host cell are of most interest. Viruses use cell surface glycans, such as heparan sulfate, sialic acid, and histo-blood group antigens, to initiate infection [6]. Antivirals that mimic these sugars can block viral entry by occupying the binding sites, blocking the virus’s natural attachment to the cell surface. Notably, heparin [7], carrageenan [8–10], and nanoparticles [11] have demonstrated antiviral activity by mimicking heparan sulfate [12]. Derivatives of sialic acid were also used in vitro against paramyxovirus [13, 14], and mimics of histo-blood group antigens have shown efficacy against norovirus [15]. Since many viruses utilize the same surface glycans, attachment inhibitors are one of the most promising strategies for developing broad-spectrum antivirals.

A significant limitation of such antivirals is their reversible interaction with the virus [6, 16]. Once administered, the antiviral may dilute within the body [6, 16]. For effective viral inhibition, such antivirals must exhibit either a high binding affinity for irreversible attachment or possess virucidal properties to neutralize the virus entirely [6, 11, 16]. Our previous work demonstrated that modified β-cyclodextrins (CD) can mimic essential cell surface glycans. CD modified on their primary face with multiple undecyls terminated with 6’sialyl-N-acetyllactosamine (6’SLN) proved to efficiently mimic sialic acid. CD modified with multiple 11-mercapto-1-undecanesulfonate (MUS) mimics heparan sulfates [17–20], making them promising antiviral agents. These macromolecules have virucidal activity without the toxicity typically associated with other virucidal compounds [17–19]. The only exception is SARS-CoV-2 where CD-MUS showed no virucidal activity [20]. However, both CDs have some limitations: in fact, CD-6’SLN exhibits greater potency in vitro compared to CD-MUS but has so far only been shown to inhibit several strains of human influenza, while did not show inhibitory activity against avian strains [17–20]. This is linked to the specificity of avian strains for α2.3 sialic acid glycans, while human strains have preference for α2.6 sialic acid glycans but can also be inhibited by α2.3 expressing molecules [18, 21].

We, therefore, aimed to identify a single strategy mimicking both heparan sulfates and sialic acid harboring a pan-respiratory virus potential. Such broad-spectrum antiviral is highly needed for pandemic preparedness, but also for known viruses for which there are no or limited approved antivirals on the market.

Here we show the results of a single modified cyclodextrin capable of targeting both heparan sulfate and sialic acid dependent viruses. This new compound showed potent efficacy against hPIV3, RSV, human and avian Influenza viruses and SARS-CoV-2. The broad-spectrum activity was confirmed in human respiratory tract tissue model and in vivo.

## Results

### Antiviral activity of modified β-cyclodextrins against hPIV3

To develop a pan-respiratory antiviral strategy which can be used against clinically relevant human respiratory viruses but also for pandemic preparedness against avian strains of influenza, a combination of previously synthesized CDs was initially considered as an approach to mimic simultaneously sialic acid and heparan sulfate. To investigate the benefit of this approach, hPIV3 was considered as a good candidate since this virus is known to depend on α2,3 sialic acid and heparan sulfate [14, 22–27]. To test antiviral activity against actively circulating strains, two clinical isolates were used in parallel with a laboratory strain (ATCC).

A glycan array assay was used to investigate hPIV3 glycan attachment. After overnight incubation of hPIV viruses at 4°C, the presence of viruses was quantified by the fluorescence intensity of the positive spots in the array. Our data showed that heparin octasaccharide and sialyl-lacto-N- neotretatose d (SLNT), an α2,3 sialic acid glycan, were common hits among the three strains tested (Figure S1). To further validate the hPIV3 dependency for sialic acid and heparan sulfates, we tested the capacity of hPIV3 to infect A549 cells deficient in the expression of sialic acid on their surface by solute carrier family 35 member A1 (SLC35A1) knockout, or LLCMK2 cells treated with sodium chlorate (Figure S2). Each strain of hPIV3 showed a significant reduction in infectivity for sialic acid knockout cells, comparable to EV-D68 used as a positive control for sialic acid dependency, and for cells with reduced sulfation of heparan sulfate proteoglycans, as for RSV used as a positive control for heparan sulfate dependency.

To mimic heparan sulfate, CD-MUS [17] (Figure 1) was used. To mimic sialic acid instead, the previously identified CD-6’SLN was not usable since it is not active against avian strains of influenza virus [18] and hPIV3 binds to α2,3 sialic acid (Figure S1) [14, 23–27]. Consequently, CD- 3’SLN [18] (Figure 1) was tested as an alternative harboring an α2,3 sialic acid glycan. However, it was not effective against hPIV3, and CD-SA [28], a CD harboring the monosaccharide sialic acid, showed low potency against HPIV3 and IAV H5N1 (Figure 1, S3A). Therefore, a new CD mimicking SLNT (CD-SLNT) was synthesized where the full SLNT glycan is grafted on the primary face of the β-cyclodextrin through an undecyl linker (Figure 1). Both CD-MUS and CD-SLNT showed no toxicity in vitro (Figure S4A, B).

**Figure 1.**
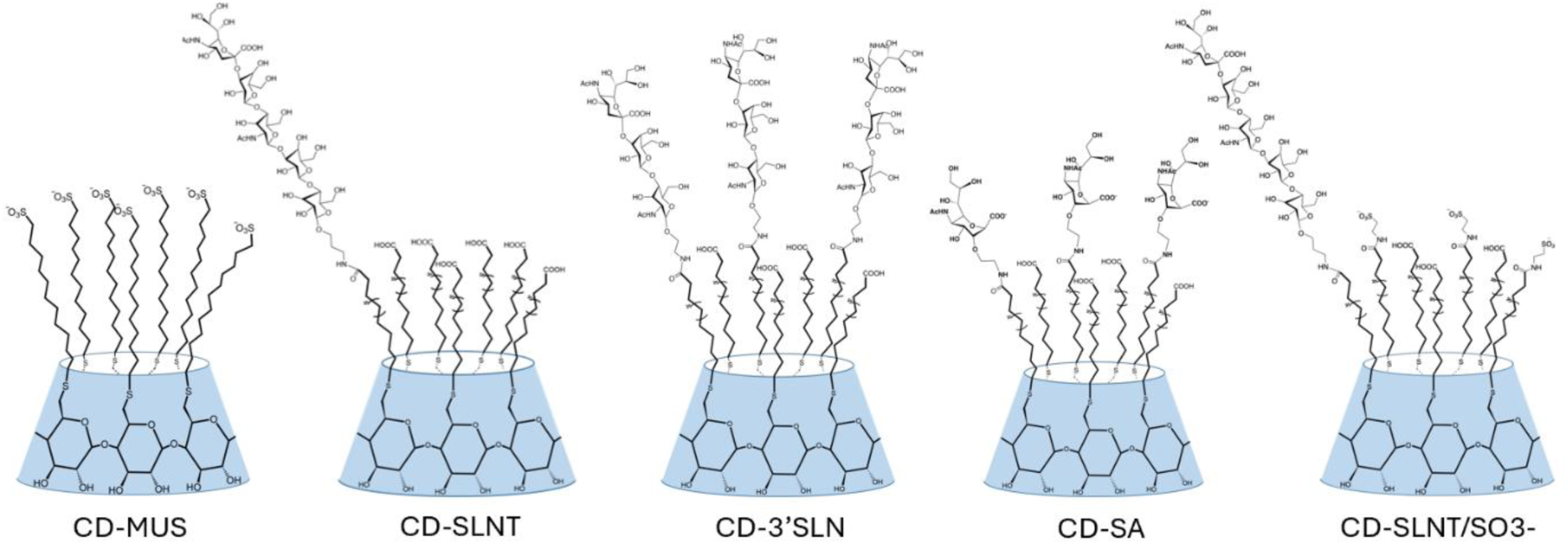
Schematic representation of modified β-cyclodextrins. Schemes of the different CD (the core is represented in blue). These drawings do not consider the possible configurations the substituents can adopt on the primary face of the cyclodextrin, but only the number of substituents.

To test the antiviral activity of these CDs, the laboratory and clinical strains of hPIV3 were incubated with CD-MUS or CD-SLNT for 1 hour at 37°C before infection. The infectivity was evaluated 3 to 5 days post-infection by plaque assay without the addition of macromolecule after infection. Additionally, the virucidal activity of these materials was assessed by incubating viruses (10^5^ pfu) and a fixed concentration of the drug (100 µg/mL CD-MUS or 300 µg/mL CD-SLNT) before serially diluting the mix and determining the viral titer by plaque assay.

CD-MUS showed an effective concentration inhibiting 50% of virus infection (EC_50_) of 2.5, 1.5, and 0.16 µg/mL (equivalent to 819.26, 496.75 and 48.98 nM) for the laboratory, clinical #1 and #2 strains respectively (Figure 2A). CD-MUS was also shown to be virucidal against the three hPIV3 strains, inducing a complete loss of infectivity of the laboratory and clinical #2 strains, and a significant 1.45 log decrease of infectivity for the clinical #1 hPIV3 isolate (Figure 2B).

**Figure 2.**
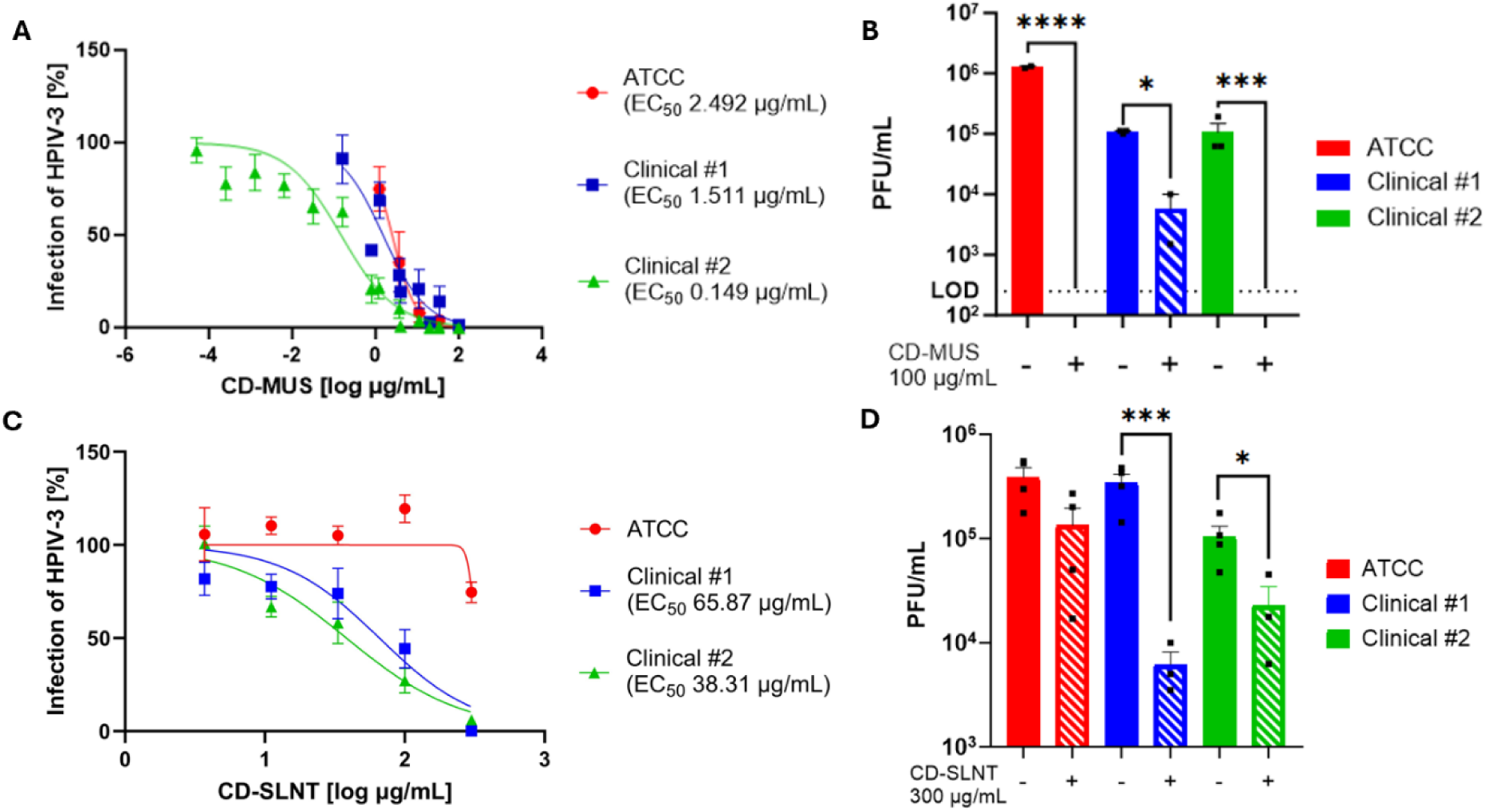
Antiviral activity of CD-MUS and CD-SLNT against hPIV3. Inhibition (A, C) and virucidal activity (B, D) of CDs. Inhibition was performed by incubating 1h at 37°C laboratory and clinical strains of hPIV3 with CD-MUS (A) or CD-SLNT (C) before infection on cells. Antiviral activity was assessed by plaque assay counted manually. Data represent mean ± SEM of two (A - ATCC), three (A – clinical #1, C), and six (A – clinical #2) independent experiments. Nonlinear regression with variable Hill slope and constraints for the bottom and top (0 and 100 respectively) were performed to compute EC_50_. (B, D) Virucidal experiments were done by incubating viruses for 1 hour with 100 µg/mL CD-MUS (B) or 2 hours with 300 µg/mL CD-SLNT (D) at 37°C. The virus-drug mix was then serially diluted. Viral titer was assessed by plaque assay counted manually. Data represent mean ± SEM of two independent (B, D) experiments. Two-tailed t-tests between untreated and treated conditions were performed. Limit Of Detection (LOD)* P< 0.0332, *** P<0.0002, **** P<0.0001

Next, the number of SLNT glycans on the CD was optimized. Each CD has the potential to have at maximum 7 active ligands grafted on the primary face. An equivalence of 1 is synthesized when the number of moles of ligands added to the chemical reaction corresponds to 7 times the quantity of moles of CD. CD-SLNT with 0.15 equivalence (i.e. corresponding to a theoretical mean of 1.05 SLNT grafted) showed the best antiviral activity with an EC_50_ of 66 and 38 µg/mL (17.99 and 10.47 µM) for clinical #1 and #2 hPIV3 respectively; however no effect was found against the laboratory strain (Figure 2C, S3A). CD-SLNT also showed virucidal activity with a 1.81 log and 0.84 log decrease in infectivity of clinical #1 and #2 isolates respectively (Figure 2D). Importantly as a control, SLNT alone did not show antiviral activity against hPIV3 clinical #2 (Figure S3A).

### Interference between CD-MUS and CD-SLNT

The combination of CD-MUS and CD-SLNT was then evaluated against hPIV3 infection. CD-MUS and CD-SLNT were mixed at different concentrations and tested against the different strains of hPIV3. Synergy scores were then obtained with SynergyFinder 3.0 [29] (Figure 3).

**Figure 3.**
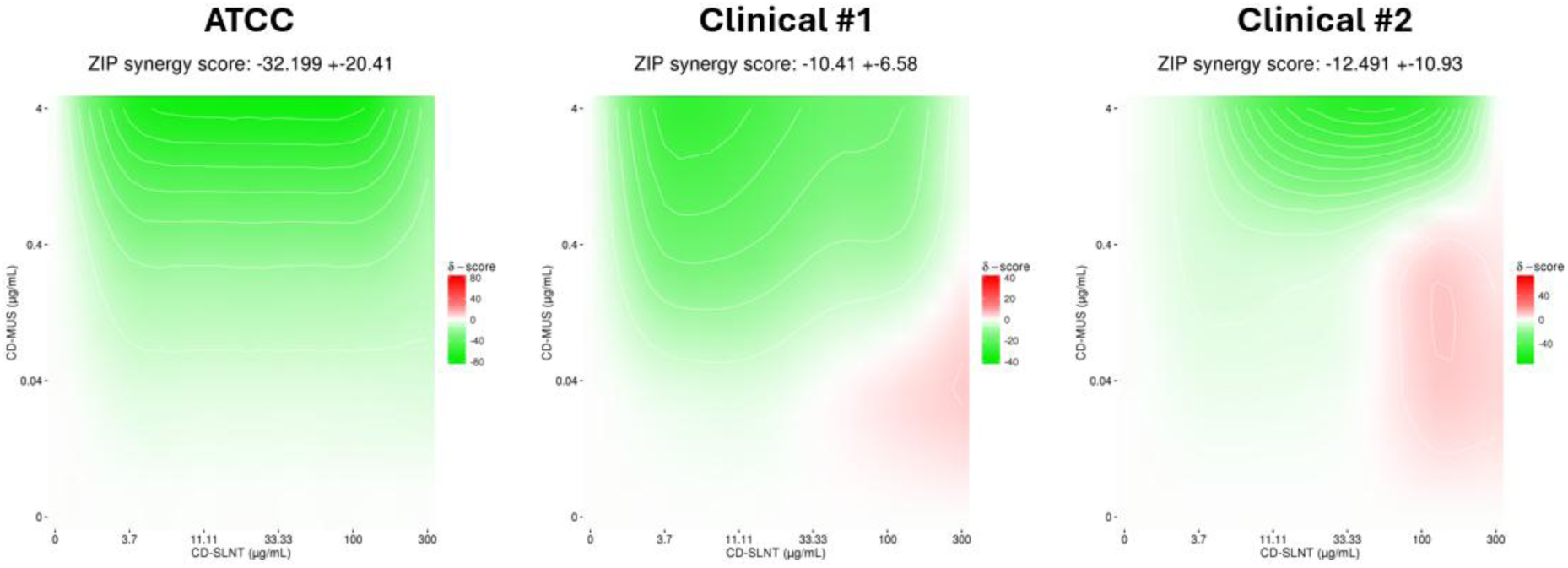
Combination assays using CD-MUS and CD-SLNT. CD-MUS and CD-SLNT were mixed at different concentrations and incubated for 1 hour at 37°C with the laboratory or the two clinical strains of hPIV3 before infection of LLCMK2 cells. Inhibition percentages were calculated by plaque assay of two independent experiments and then evaluated with SynergyFinder 3.0 [29]. The reference algorithms (HSA, Loewe, Bliss, and ZIP) were applied. Green, white, and red areas indicate antagonistic, additive, and synergistic areas respectively.

For the laboratory hPIV3 strain, results showed a clear antagonism between CD-MUS and CD- SLNT. On the opposite, for hPIV3 clinical isolates, synergy scores are at the limit between antagonism and additivity (i.e. -10). All the different scores to evaluate synergism (Highest single agent (HAS), Loewe, Bliss, or Zero interaction potency (ZIP) [30–35]), were concordant with the antagonistic effect (Table S1).

This antagonist effect was investigated by a computational approach. SLNT was docked on site I of the sialic acid pocket previously described in the hemagglutinin-neuraminidase (HN) protein [36–38] (PDB 5B2D [39]) (Figure 4A). The sialic acid was stabilized by a complex of three arginines (Figure 4B, S5A). SLNT was not possible to dock on site II at the dimer interface [36, 40, 41]. Molecular dynamics of 500 ns were performed with 10 CD-MUS around the hemagglutinin-neuraminidase protein (PDB 4MZA [42]). Most of the amino acids with which CD-MUS interacts by hydrogen bonds, water bridges, or ionic interactions are lysines (30.7%), asparagines (19.5%), threonines (16.5%) and arginines (12.7%) due to their polarity. Although they showed to bind mainly distal locations to the sialic acid binding site (Figure S5B), interactions were observed in and around the site I sialic acid binding pocket, including with the arginines interacting with SLNT (Figure 4C). These data support the antagonism observed experimentally since the binding of one CD could prevent the interaction of the other.

**Figure 4.**
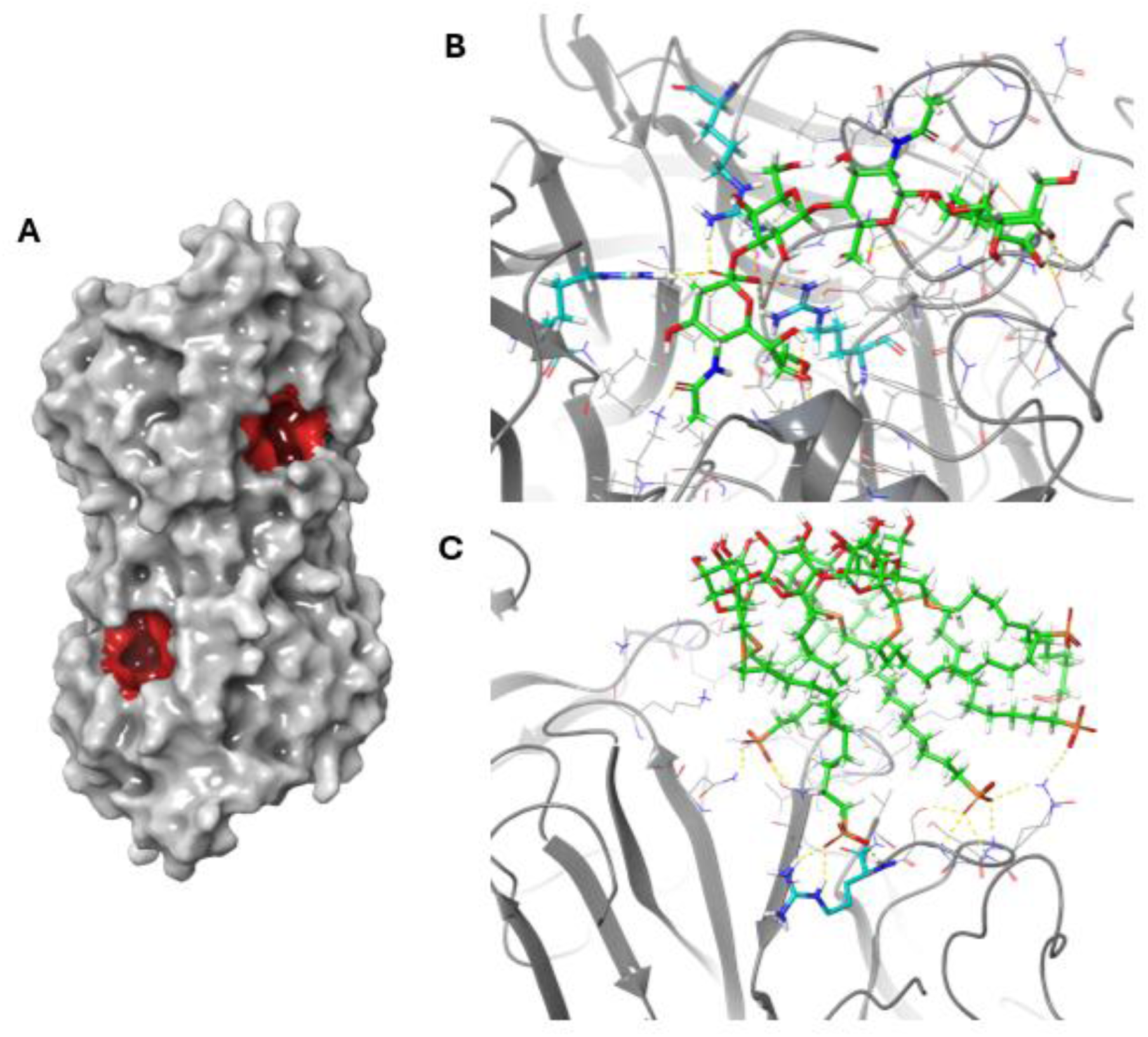
CD interactions with the sialic acid binding pocket. The two identical sialic acid binding pockets (Site I) are highlighted in red on the hemagglutinin-neuraminidase of hPIV3 (A). SLNT was docked on the boxed sialic acid binding pocket (PDB 5B2D [39]). (B) CD-MUS interactions with the HN protein were simulated by molecular dynamics and showed interaction with this pocket (PDB 4MZA [42]) (C). Ligands carbons are in green and key arginine residues are in cyan.

### Pan- activity of the dual active molecule against respiratory viruses in vitro

To overcome the antagonism of the two cyclodextrins, a new CD with SLNT and sulfonate as active epitopes on the primary face was synthesized: CD-SLNT/SO3- (Figure 1). CD-SLNT/SO3- showed lower EC_50_ on clinical isolates of hPIV3 if compared to CD-SLNT (Table 1). Additionally, molecular dynamics support the biological results, confirming that SLNT can bind to the sialic acid pocket while the sulfate can interact with other residues without interference (Figure S5C).

**Table 1.** Antiviral activity of CD-MUS, CD-SLNT, and CD-SLNT/SO3-. Effective concentration inhibiting 50% of virus infection (EC_50_). Molarities were calculated considering the main synthesized macromolecule. n.a: not assessed. † Viruses for which CD-SLNT/SO3- is more potent than both CD-MUS and CD-SLNT.

| Virus | CD-MUS (EC<sub>50</sub>)<br>[μg/mL] | CD-SLNT (EC<sub>50</sub>)<br>[μg/mL] | CD-SLNT/SO3-<br>(EC<sub>50</sub>) [μg/mL] | Virucidal<br>CD-SLNT/SO3- |
| --- | --- | --- | --- | --- |
| **hPIV3 ATCC** | 2.49<br>(819.26 nM) | >300 | 56.94<br>(14'314.23 nM) | Y |
| **hPIV3 Clinical #1†** | 1.51<br>(496.75 nM) | 65.87<br>(17'999.73 nM) | 0.19<br>(47.76 nM) | Y |
| **hPIV3 Clinical #2** | 0.15<br>(48.98 nM) | 38.31<br>(10'468.64 nM) | 0.62<br>(154.61 nM) | Y |
| **RSV** | 0.74<br>(242.62 nM) | >300 | 2.29<br>(576.69 nM) | Y |
| **SARS-CoV-2†** | 5.56<br>(1'827.89 nM) | >300 | 0.35<br>(86.73 nM) | Y |
| **IAV H1N1†** | 6.11<br>(2'007.39 nM) | 0.13<br>(36.32 nM) | 0.052<br>(13.07 nM) | Y |
| **IBV** | 20.09<br>(6'604.73 nM) | 2.61<br>(711.85 nM) | 4.37<br>(1.09 μM) | Y |
| **IAV H5N1** | 54.86<br>(18'035.61 nM) | 0.15<br>(39.62 nM) | 0.57<br>(143.04 nM) | Y |
| **IAV H7N1** | n.a | n.a | 1.39<br>(350.44 nM) | n.a |
| **HSV-2** | 1.11<br>(364.92 nM) | >300 | 20.14<br>(5'063.02 nM) | n.a |

More importantly, this dual-active molecule can mimic simultaneously heparan sulfate and sialic acid and have a broader antiviral activity. CD-SLNT/SO3- was therefore tested against a panel of viral pathogens. The macromolecule showed activity against IAV (H1N1), IBV, SARS-CoV-2, RSV, and HSV-2 and also against two different avian strains of IAV (H5N1 and H7N1) (Table 1) in the low micromolar to nanomolar range. Virucidal activity of CD-SLNT/SO3- was confirmed against hPIV3, RSV, Influenza viruses, and importantly against SARS-CoV-2 as well (Table 1, Figure S6). Importantly, the investigation of the virucidal kinetics on Influenza viruses, revealed significant inactivation even without preincubation between the virus and the compound (Figure S7).

Characterization of CD-SLNT/SO3- by high-performance liquid chromatography coupled with mass spectrometry showed a mixture of CDs (Figure S8) with a prevalent species being a CD harboring one SLNT and multiple sulfonate groups. Different batches were synthesized to assess the reproducibility of the synthesis and the different batches showed consistent activity against IAV H1N1 and RSV (Figure S3C, E). Furthermore, for viruses dependent on sialic acid (hPIV3, IAV), an increase of SLNT grafted at the surface of CD-SLNT/SO3- reduced the potency of the macromolecule as also shown for CD-SLNT against hPIV3 (Figure S3B, D, E) while the effect was not observed for RSV which does not depend on sialic acid (Figure S3C).

### Resistance to CD-SLNT/SO3-

To investigate the barrier to resistance of CD-SLNT/SO3-, hPIV3 clinical #2 and IAV H1N1 were passaged 10 times in the presence of an increasing concentration of the macromolecule. After the last passage, the antiviral activity of CD-SLNT/SO3- was assessed against the virus passaged 10 times in the presence or absence of the macromolecule. After 10 passages, no resistance of hPIV3 to the macromolecule was detected (Figure 5A). However, regarding IAV H1N1, the antiviral potency of CD-SLNT/SO3- was reduced by 28 times against treated viruses compared to the untreated virus for which the macromolecule already lost 10 times its antiviral activity after 10 passages (Figure 5B). Two CD-SLNT/SO3- resistant IAV variants were purified by plaque assay and sequenced to determine whether specific mutations were present (Figure 5B). We could indeed identify several mutations in the hemagglutinin and neuraminidase genes (Table S2), two of which, K154E and G155E, are located close to the sialic acid binding site of the hemagglutinin (Figure 5C). In the GISAID database, they are observed to be present in less than 1% of the sequenced influenza viruses (Table S2).

**Figure 5.**
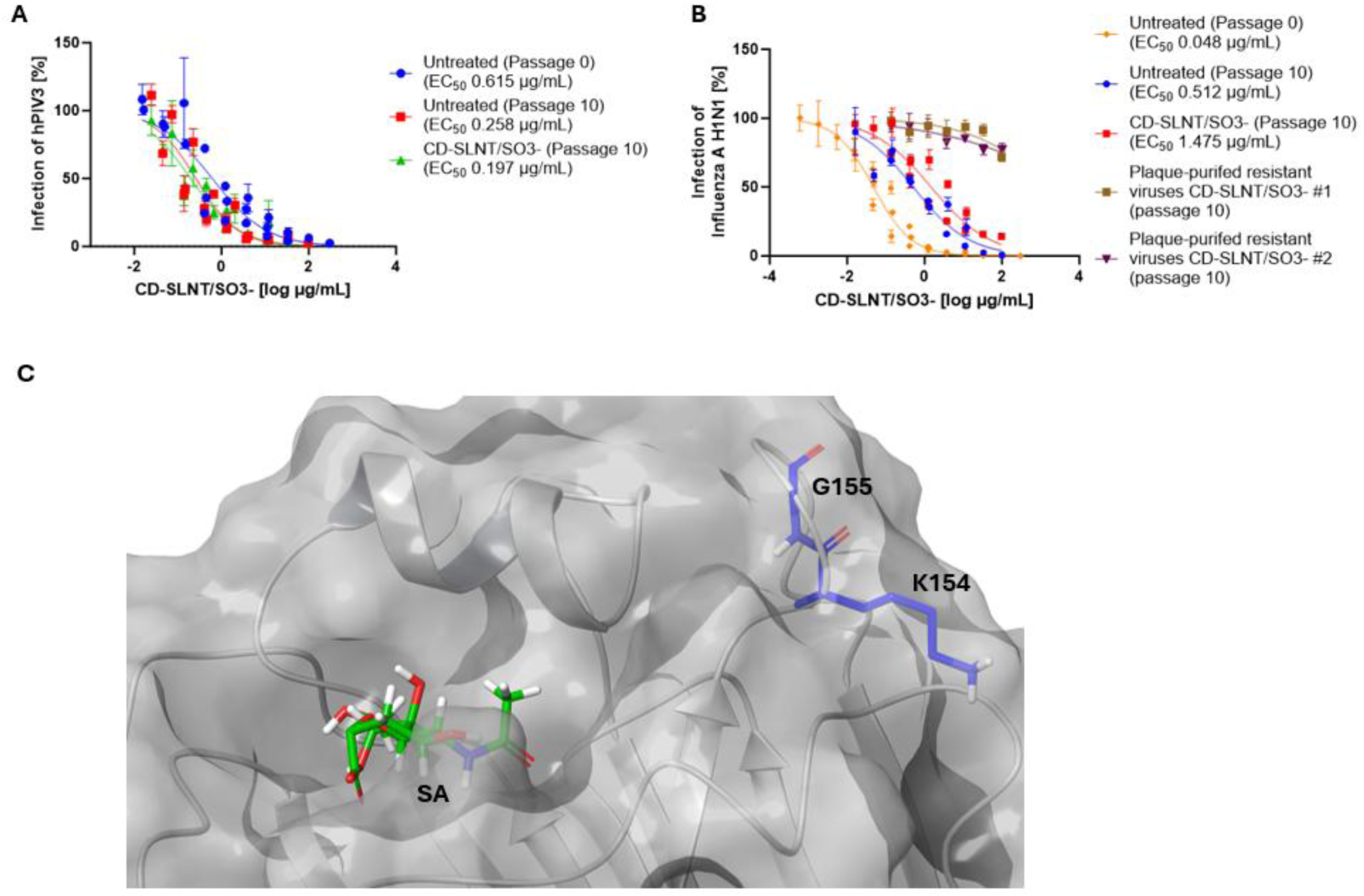
Resistance to CD-SLNT/SO3-. HPIV3 (A) and Influenza A H1N1 (A) viruses were passaged 10 times in the presence of an increasing concentration of CD-SLNT/SO3-. At the final passage, the antiviral activity of CD-SLNT/SO3- against the untreated and treated viruses was assessed. Resistant plaque-purified Influenza A H1N1 viruses were obtained (B). Data represent mean ± SEM of two (B-Plaque purified resistant CD-SLNT/SO3- passage 10), three (B-Untreated passage 10, CD-SLNT/SO3- passage 10) or four (A) independent experiments. Each independent experiment with untreated or treated virus at passage 10 was carried out using the two independent virus replicates. Nonlinear regression with variable Hill slope and constraints for the bottom and top (0 and 100 respectively) were performed to compute EC_50_. (C) Mutations observed in resistant viruses present in the hemagglutinin protein (PDB 4JTV [50]) and sialic acid (SA) at the glycan binding region are highlighted in blue and green respectively.

### Broad-spectrum activity of CD-SLNT/SO3- ex vivo

The efficacy of CD-SLNT/SO3- was then evaluated in a more relevant model for viral infections, human derived upper respiratory tract pseudostratified airway epithelium. HPIV3, SARS-CoV-2, RSV, or IAV H1N1 were pre-incubated for 1 hour at 37°C with 300 µg/mL of the dual active CD (75.42 µM). Human respiratory airways were then infected and maintained at 33°C for 4 days without further addition of the macromolecule. Viruses were collected every day by an apical wash and then quantified by RT-qPCR. In ex vivo human airways infected with viruses pre-incubated with CD-SLNT/SO3-, a significant reduction of apically released viruses was observed (Figure 6). CD-SLNT/SO3- did not show ex vivo toxicity by the measurement of cell viability or LDH release even with daily addition of the macromolecule (30 µg – 7.54 nmol) (Figure S4C, D).

**Figure 6.**
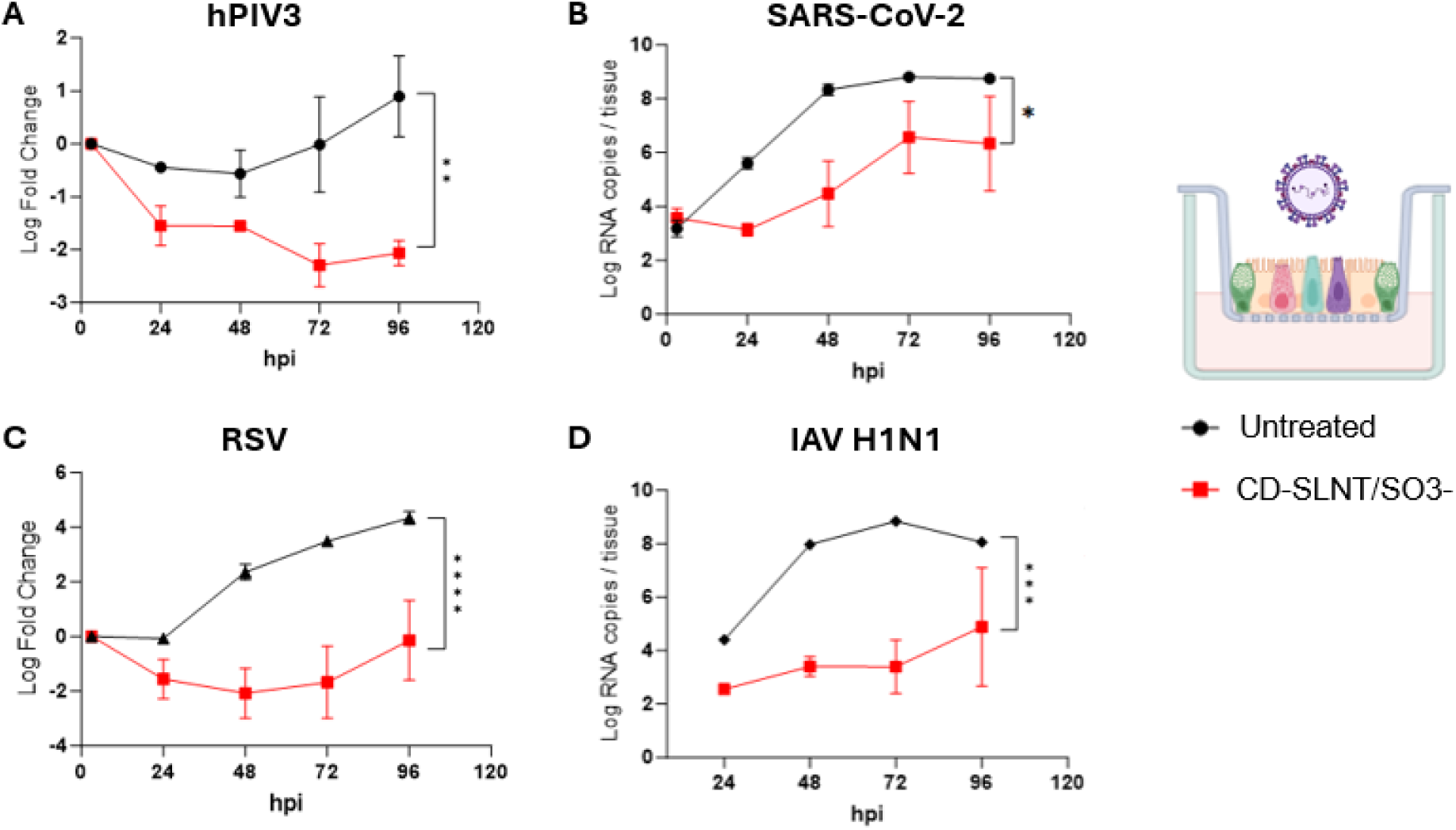
Ex vivo activity of CD-SLNT/SO3-. We incubated hPIV3 (A), SARS-CoV-2 (B), RSV (C), or Influenza A H1N1 (D) with 300 µg/mL of CD-SLNT/SO3- for 1 hour at 37°C. Human upper respiratory tract models (n = 2 (A, C, D) or 6 (B) per group) were then infected at 33°C. A daily apical wash was performed and the level of viruses released was quantified by RT-qPCR. Data represent mean ± SEM. Area under the curve followed by a two-tailed t-test was performed. ** P<0.0021, *** P<0.0002, **** P<0.0001. Cartoon created with biorender.com

### Activity in vivo of CD-SLNT/SO3-

The antiviral activity of CD-SLNT/SO3- was assessed against IAV H1N1 in a zebrafish larvae model. This model was previously shown to be relevant for influenza infection [43] and known antiviral molecules such as baloxavir acid are able to inhibit virus replication thus showing the model is suitable to test new antivirals (Figure S9). IAV H1N1 together with 100 µg/mL CD- SLNT/SO3- were co-injected into the swim bladder of zebrafish larvae without further addition of the macromolecule (Figure 7A). A pool of 10 zebrafish were harvested at different time points to quantify the amount of viral RNA. The addition of CD-SLNT/SO3- during infection inhibited the replication of IAV H1N1 virus by 1.68 log (Figure 7B). No signs of toxicity or locomotor impairment [44] were observed by the addition of CD-SLNT/SO3-, in comparison to DMSO-treated control.

**Figure 7.**
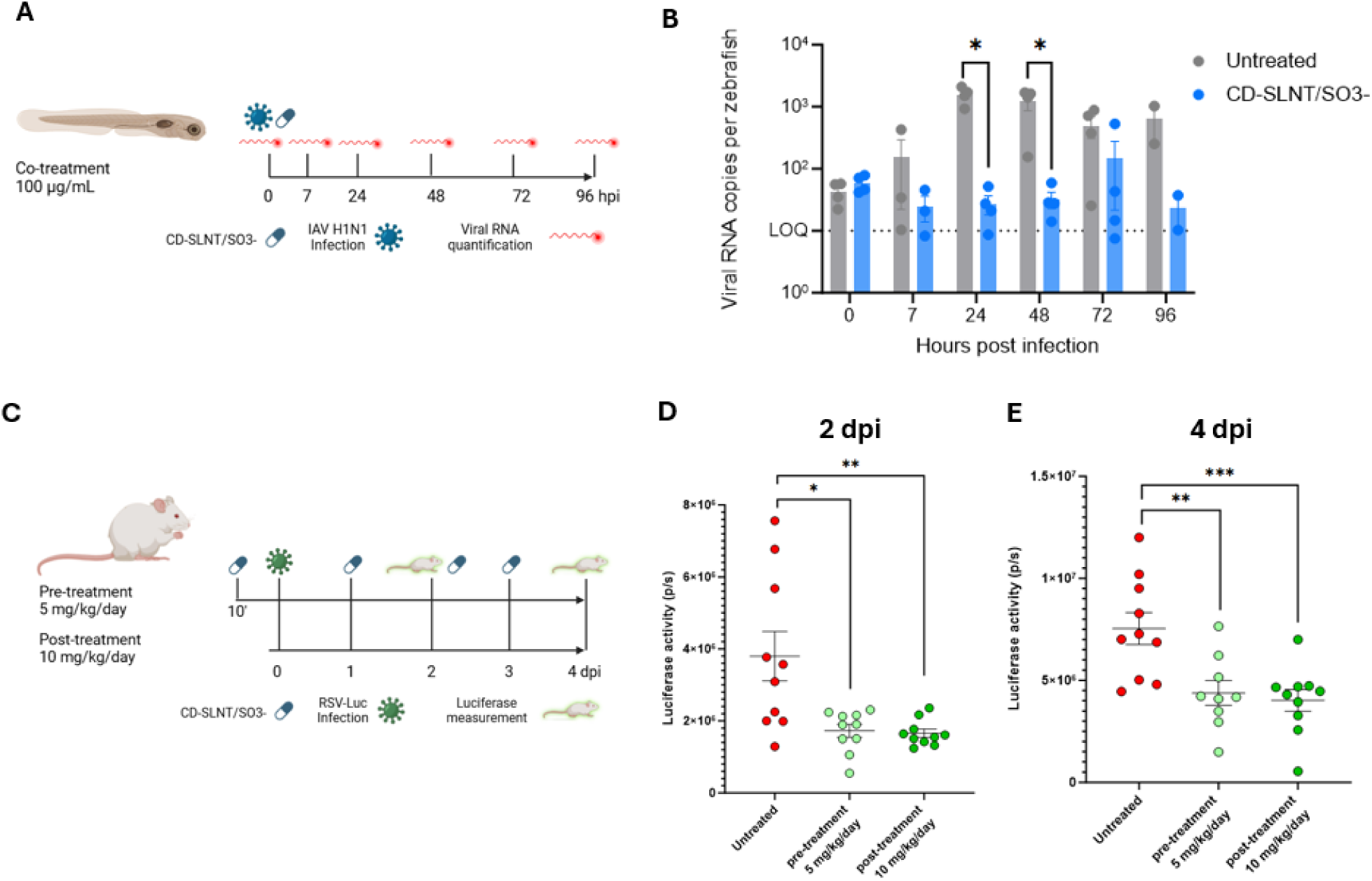
In vivo activity of CD-SLNT/SO3-. (A, B) Zebrafish larvae were infected with IAV H1N1. CD-SLNT/SO3- was co-injected at the same time at a concentration of 100 µg/mL. At 7 hpi and each day, 10 larvae are lysed for viral RNA quantification by RT-qPCR. Data represents mean ± SEM of four independent experiments. (C, D, E) Mice (n = 10 per group) were infected with RSV. (C) Mice were either treated with 5 mg/kg of CD-SLNT/SO3- 10 minutes before infection (pre-treatment) with a daily dose until 4 days post-infection (dpi) or with 10 mg/kg of CD-SLNT/SO3- starting 1 dpi (post-treatment). (D, E) RSV replication was assessed by luciferase activity at 2 (D) and 4 (E) dpi. Data represents mean ± SEM of a single experiment. (Two-tailed Mann-Whitney tests were performed to compare untreated and treated conditions. * P< 0.0332, ** P<0.0021, *** P<0.0002. Schematic view created with biorender.com

The antiviral activity of CD-SLNT/SO3- was further evaluated against RSV in vivo in a mouse model. Mice were treated with 5 mg/kg/day of CD-SLNT/SO3- 10 min before infection with RSV and daily for 3 days (Figure 7C). The luciferase activity in the nose and lungs of mice was measured at days 2 and 4 post-infection. CD-SLNT/SO3- treatment was shown to inhibit RSV replication at days 2 and 4 post-infection (Figure 7D, E). These results were reproduced in an independent experiment (Figure S10A). No loss of weight was observed by the administration of CD-SLNT/SO3- in mice (Figure S10B).

Finally, the therapeutical activity of CD-SLNT/SO3- was also evaluated by treating RSV-infected mice starting 1 day post-infection with 10 mg/kg/day of CD-SLNT/SO3- (Figure 7C). In this setting, a comparable global reduction in viral replication was observed (Figure 7D, E). For both types of administration, a significant 0.3 log reduction of infection in the lungs of treated mice was observed (Figure S10C) and in the nose of pre-treated mice at day 4 post-infection (Figure S10D).

## Discussion

Respiratory viruses continue to pose a dual challenge: a significant clinical burden and a persistent pandemic threat. These infections affect individual lives and the global economy, with vulnerable populations—such as children, immunocompromised individuals, and the elderly—often exhibiting insufficient immune responses [45, 46]. For these groups, and for viruses without effective vaccines, antivirals remain critically needed. Moreover, the risk of zoonosis is ever-present, as demonstrated by the scrutiny of avian strains of influenza for their zoonotic potential. While vaccination is the preferred strategy for controlling known pathogens, the unpredictable emergence of novel viruses amplifies the need for broad-spectrum antivirals. Our previous research focused on CD to mimic glycans such as sialic acid and heparan sulfate [11, 17, 19, 20, 47]. In this study, we made a breakthrough achievement by designing and evaluating the antiviral activity of a dual-active virucidal macromolecule that mimics both sialic acid and heparan sulfate, resulting in a single macromolecule able to inhibit major respiratory viral pathogens and avian strains of influenza virus.

An initial combination of the sulfonated CD-MUS and the sialylated CD-SLNT was explored against hPIV3, a virus known to use sialic acid and heparan sulfate as attachment receptors (Figure S1, S2) [36, 48]. Both macromolecules showed antiviral and virucidal activity independently against hPIV3 due to their structural features (Figure 2) [17, 18]. However, the combination of both CDs resulted in antagonism (Figure 3), which was explored with an in silico approach, showing the potential interference of the two macromolecules by binding to the residues close or within the sialic acid binding region (Figure 4, S5A, B). A similar antagonism is expected as well for other viruses such as IAV H1N1. Indeed, lysine, asparagine, threonine, and arginine, the key interacting residues with CD-MUS (Figure S5B), are locally present in the sialic acid binding region on the hemagglutinin of IAV [49–51] and could interact with the glycans mediating the interference.

We therefore designed and synthesized a dual-active CD that incorporates SLNT and sulfonate groups on its primary face, referred to as CD-SLNT/SO3-, providing a unique strategy to simultaneously target viruses using either or both receptors (Figure 1). Indeed, the macromolecule was inhibiting and had virucidal activity against major respiratory viruses (IAV H1N1, IBV, SARS-CoV-2, RSV, and hPIV3) and against two avian strains of influenza virus (IAV H5N1 and IAV H7N1) (Table 1, Figure S6). While CD-SLNT/SO3- showed improved antiviral efficacy compared to either CD-MUS or CD-SLNT for each virus tested, the dual-active material demonstrated superior potency against a clinical isolate of hPIV3, IAV H1N1, and SARS-CoV-2 compared to both single active CD (Table 1) suggesting that the mixture of ligands confers to the material new chemical properties. Indeed, the dual-active macromolecule has better efficacy than previous attachment inhibitors. For example, CD-SLNT/SO3- showed nanomolar activity against SARS-CoV-2 and IAV H1N1 while other virucidal attachment inhibitors designed with a similar approach but with a different scaffold than β-cyclodextrin resulted in a micromolar activity [52]. CD-SLNT/SO3- has virucidal activity against SARS-CoV-2 in contrast to CD-MUS (Figure S6B) [20]. Furthermore, the kinetics of virucidal activity observed against Influenza viruses suggested a faster inactivation by CD- SLNT/SO3- (Figure S7) than previous CD developed [17].

This broad-spectrum efficacy not only establishes CD-SLNT/SO3- as a versatile antiviral candidate but also underscores its transformative potential for clinical disease management. By targeting viruses that utilize either sialic acid or heparan sulfate, this macromolecule could address respiratory viruses not yet tested, such as hPIV1 known to have similar sialic acid glycan affinity than hPIV3 [23, 53], human metapneumovirus, human coronavirus NL63, and SARS-CoV known to use heparan sulfate as attachment receptor [54–57], IAV H7N9 and virtually all other avian viruses which are reported to bind α2,3 sialic acid [58, 59]. Importantly new emerging sarbecoviruses or new strains of avian influenza virus could also be inhibited by the CD, making it an important tool for pandemic preparedness. In a clinical setting, such a molecule could revolutionize disease management by providing a single, broad-spectrum treatment option. This would reduce reliance on pathogen-specific antivirals with a narrow spectrum activity. Incorporating CD-SLNT/SO3- into therapeutic strategies could facilitate rapid responses during outbreaks and improve outcomes for high-risk populations, including those for whom vaccines are ineffective. While further in vivo testing is needed, the scope of activity achieved here already represents a major breakthrough, underscoring the imperative to continue developing such game-changing molecules.

Despite the promising results, the synthesis of the macromolecule presents challenges, as it produces a mixture of CDs which represents the main limitation of our study. However, various batches of CD-SLNT/SO3- tested against IAV H1N1 and RSV revealed consistent antiviral activity (Figure S3C, E). Characterization via HPLC-MS indicated that the most abundant species is CD with one single SLNT and multiple sulfonate groups (Figure S8), this is in line as well with the reduced activity of batches with higher degrees of substitution (Figure S3B, D, E). These findings also highlight that developing multivalent materials does not necessarily lead to higher potency.

The barrier to resistance of CD-SLNT/SO3- was evaluated against hPIV3 and IAV H1N1. While for hPIV3, no signs of resistance were observed (Figure 5A), IAV H1N1 revealed a reduction in potency after ten passages (Figure 5B). Although cell adaptation might already alter the efficacy of CD-SLNT/SO3-, additional mutations were identified on two different resistant viruses that can abolish the antiviral activity of the macromolecule (Table S2): K154E and G155E mutations are located near the sialic acid binding site (Figure 5C), validating the mechanism of action. Notably, K154E has been shown to decrease potency against CD-SA after passages with a combination of interferon and CD-SA [28], while G155E might increase α2,6 sialic acid preference [60, 61]. A possible explanation is that SLNT, which terminates with an α2,3 sialic acid, would bind less effectively in the altered sialic acid binding pocket, as the virus could evolve to use preferentially α2,6 sialic acid. Despite this potential risk of resistance, analysis of over 35,000 sequences from GISAID indicated that the nucleotide mutations responsible for K154E and G155E were present in less than 1% of sequences (Table S2). This suggests that the likelihood of encountering a virus in clinical settings harboring these mutations is low. However, resistance could emerge during infection, and considering this possibility, combination therapy should be considered to delay resistance. CD-SLNT/SO3- could be associated with other CDs previously described such as CD- 6’SLN or another antiviral, DAS181, which has a host-directed activity by cutting sialic acid and finished a Phase II clinical study against Influenza infection [18, 62].

The activity of CD-SLNT/SO3- was verified in two more complex models. In ex vivo human respiratory tissues, which proved to be relevant for respiratory viruses [63–66], the macromolecule reduced the level of apically released hPIV3, RSV, IAV H1N1, and SARS-CoV-2 (Figure 6). The broad-spectrum efficacy of CD-SLNT/SO3- was then confirmed in zebrafish larvae against IAV H1N1 and in mice against RSV. Administration at the time of infection showed efficacy in both models (Figure 7). Additionally, in mice the use of therapeutic administration against RSV showed similar efficacy further confirming the antiviral activity in vivo (Figure 7B, C, S10).

In conclusion, we have presented here what, to the best of our knowledge, is the first non-toxic virucidal pan-respiratory antiviral. All the findings presented support further investigation of CD- SLNT/SO3- as a potential treatment for human respiratory viral infections effective as well for avian strains of influenza virus which could be formulated, for example, as an inhalable dry powder [67], with a high likelihood of being effective against newly emerging respiratory viruses, making it key for pandemic preparedness.

## Materials and Methods

### Cell and viruses

LLCMK2 (ATCC CCL-7), A549 (ATCC CRM-CCL-185), MDCK-Siat, A549 SLC35A1 KO, and Vero E6 (ATCC CRL-1586) were kindly given by Prof. Caroline Tapparel, Prof. Mirco Schmolke, and Prof. Gary Kobinger. Cells were maintained in Dulbecco’s Modified Eagle Medium (DMEM) (Gibco) with 10% Fetal Bovine Serum (FBS) (Pan Biotech) and 1% penicillin/streptomycin (Gibco) at 37°C and 5% CO2.

HPIV3 was purchased on ATCC and two clinical samples were isolated from clinical specimens. IAV (A/Vietnam/1203/2004 (H5N1), and A/Netherlands/602/2009 (H1N1)) were provided by Prof. Mirco Schmolken and Influenza B virus (IBV) (B/Washington/02/2019) by the Sentinella Center for Influenza Virus at the University Hospital of Geneva. Enterovirus D68 (EV-D68) was given by Prof. Caroline Tapparel. SARS-CoV-2 BA.1 Omicron was isolated from a clinical sample [68]. RSV-GFP was obtained from [69].

### Glycan Array

The glycan array was purchased from ZBiotech (ref 10601-8S) harboring 100 different glycans at its surface. The array was initially blocked for 1 hour at room temperature with Phosphate-buffered saline (PBS) (Bischel), 1% Bovine Serum Albumin (BSA) (AppliChem), and 0.05% Tween-20 (Sigma) (blocking solution). Non-diluted viral stock of hPIV3 (laboratory and clinical isolates) (ranging from 1.69 to 4.79 10^4^ focus forming unit (ffu)/mL) was added to two subarrays for each strain overnight at 4°C. The array was then washed three times with PBS, 0.05% Tween 20 (wash solution) before fixation with PBS, 4% formaldehyde. The glycan array was incubated in the presence of anti-parainfluenza (1:10) (Light Diagnostics) in blocking solution for 1 hour at 37°C and then washed three times for 5 minutes each. The array was then incubated with anti-mouse AlexaFluor 488 (1:1500) (Invitrogen) in the blocking solution for 1 hour at 37°C. After three washes of 5 minutes, the array was disassembled and immersed in the wash solution for 10 minutes. Deionized water was used for the last wash for 5 minutes before drying the glycan array. The array was visualized with Leica DMi 8 and the intensities of the different positive spots were quantified with ImageJ.

Sialic acid Knock out cells’ infection assay A549 and A549 SLC35A1 KO cells (16’000 cells/well) were seeded in a 96-well plate. Cells were infected with hPIV3 or EV-D68 with a 1:10 dilution of the viral stocks (ranging from 1.69 to 4.79 10^4^ ffu/mL for hPIV3 and 1.48 10^7^ ffu/mL for EV-D68) in DMEM, 200 ng/mL TPCK Trypsin (Sigma) or DMEM, 2.5% FBS respectively. After 1 hour at 37°C, infected cells were incubated in the infection medium for 1 day at the same temperature. After fixation with pure methanol, cells were incubated in the presence of PBS, 1% BSA, and 0.05% Tween-20 for 20 minutes at room temperature. Cells were then incubated with either anti-parainfluenza (1:10) or anti-enterovirus (1:500) in PBS, 1% BSA, and 0.05% Tween-20 for 1 hour at 37°C, followed by three washes with PBS, 0.05% Tween 20. Cells were then incubated with PBS, 1% BSA, and 0.05% Tween-20, anti-mouse horseradish peroxidase (HRP) (1:1000) (Cell Signaling) for 1 hour at 37°C. After three washes, tetramethyl benzidine (Invitrogen) was added to react with HRP which was stopped by the addition of HCl after enough differentiation of the signal compared to the background. Absorbance was read at 450nm with a Microplate reader.

### Heparan sulfate dependency experiment

LLCMK2 cells were passed at least five times in the presence of 30 mM sodium chlorate (NaClO3) to inhibit sulfation of heparan sulfate proteoglycans. LLCMK2 cells passed similarly without NaClO3 were used for the untreated condition.

For hPIV3 infection, cells were seeded in a 24-well plate (80’000 cells/well). One hundred plaque-forming units (pfu) of the laboratory or clinical strains were used to infect cells for 1 hour at 37°C in DMEM, 200 ng/mL TPCK Trypsin. An overlay of DMEM, 0.5% Methylcellulose (MTC) (Sigma), 200 ng/mL TPCK Trypsin with or without 30 mM NaClO3 was added to the cells. Infected cells were incubated for 3 (laboratory strain) to 5 (clinical strains) days at 37°C. A solution of water, 20% ethanol, 0.1% crystal violet (Sigma) was used to fix and stain cells. Plaques were quantified manually.

For RSV-GFP infection, cells were seeded in a 96-well plate (10’000 cells/well). Two hundred ffu of RSV-GFP was used to infect cells for 1 hour at 37°C in DMEM 2.5% FBS. DMEM, 2.5% FBS with or without 30 mM NaClO3 was added to the cells. Infected cells were incubated for 1 day at 37°C. Cells expressing GFP were then counted manually to measure infectivity.

### Synthesis and characterization of modified β-cyclodextrin

CD-MUS, CD-3’SLN, CD-6’SLN, and CD-SA were synthesized similarly to previously published protocols [17, 18, 28]. All other CD used the synthesis of β-cyclodextrin with undecyl chains grafted on the primary face with carboxy groups activated with N-Hydroxysuccinimide (NHS), ready for conjugation with a primary amine [47, 67]: for details on the synthesis, consult the supplementary information (SI). Generally, the aminated end group was reacted to a specific proportion of NHS activated β-cyclodextrin in dimethyl sulfoxide (DMSO) (Sigma) in the presence of triethylamine (TEA) followed directly by dialysis against MilliQ water using 1.5 or 2 kDa regenerated cellulose membranes (Spectrum laboratories). The contents of the dialysis bag were then freeze-dried and collected for use.

Batches of CD-SLNT: Activated β-cyclodextrin was reacted with different ratios of α2,3-sialyl Lacto-N-neotetraose aminopropylglycoside (Neu5Acα2-3Galβ1–4GlcNAcβ1–3Galβ1–4Glcβ1-4 propylamine SLNT – Asparia Glycomics). For initial tests, batches with 1.2, 0.6, 0.3, and 0.15 equivalents of SLNT per NHS group were synthesized and tested against the viruses. The reactions using only SLNT were done as follows: using a TEA (Sigma) stock solution of 70 µL TEA in 500 µL DMSO (i) 7mg (0.0021 mmol) of activated β-cyclodextrin mixed with 0.3 equivalents of SLNT (4.75 mg, 0.0045 mmol) in 5 mL of DMSO to which 20 µL of the TEA solution was added into a 5 mL round bottom flask and magnetically stirred for 12 h, followed by dialysis against milliQ (ii) 7 mg (0.0021 mmol) of activated β-cyclodextrin mixed with 0.6 equivalents of SLNT (9.5 mg, 0.009 mmol) in 5 mL of DMSO to which 20 µL of the TEA solution was added into a 5 mL round bottom flask and magnetically stirred for 12 hours followed by dialysis against milliQ (iii) 7 mg (0.0021 mmol) of activated β-cyclodextrin mixed with 1.2 equivalents of SLNT (19 mg, 0.018 mmol) in 5 mL of DMSO to which 20 µL of the TEA solution was added into a 5 mL round bottom flask and magnetically stirred for 12 hours followed by dialysis against milliQ. All batches were freeze-dried and collected as a white powder.

CD-SLNT/SO3-: (i) batch #1: To 50 mg (0.0150 mmol) of activated β-cyclodextrin, 0.15 equivalents of SLNT was used per NHS group (18.7 mg, 0.0177 mmol) with 8 mg (0.064 mmol) of taurine (Sigma) in 10 mL of DMSO. The mixture was magnetically stirred for 12 h, dialyzed, and freeze-dried. (ii) batch #2 and #3: 200 mg of activated β-cyclodextrin (0.06 mmol) with 74.8 mg of SLNT (0.07 mmol) and 32 mg of taurine (0.255 mmol) were added to 20 mL of DMSO and stirred magnetically for 12h, then dialyzed and freeze-dried. (iii) A batch with 2 SLNTs (0.28 equivalence) and 5 taurines per CD – 50 mg of activated β-cyclodextrin (0.0150 mmol) with 31.68 mg of SLNT (0.03 mmol) and 9.38 mg of taurine (0.075 mmol) with 50 µL of the TEA solution were stirred magnetically in 10 mL DMSO for 12h, dialyzed and freeze-dried.

CD-3’SLN/SO3-: 50 mg of activated β-cyclodextrin (0.015 mmol), 21.53 mg of 3’sialyl-N-acetyllactosamine (3’SLN) (0.03 mmol), and 9.38 mg of taurine (0.075 mmol) were mixed in 10 mL of DMSO and stirred magnetically for 12h, followed by dialysis and freeze-drying.

CD-6’SLN/SO3-: 50 mg of activated β-cyclodextrin (0.015 mmol), 21.53 mg of 6’’sialyl-N-acetyllactosamine (6’SLNEtNH2) (0.03 mmol), and 9.4 mg of taurine (0.075 mmol) were mixed in 10 mL of DMSO and stirred magnetically for 12h, followed by dialysis and freeze-drying.

CD-SA/SO3-: 50 mg of activated β-cyclodextrin (0.015 mmol) with 10.5 mg (0.03 mmol) of SAEtNH2 and 3.75 mg of taurine (0.03 mmol) were mixed in 10 mL of DMSO and stirred magnetically for 12h, followed by dialysis and freeze-drying.

CD-SO3-: 50 mg of activated β-cyclodextrin (0.015 mmol) and 10 mg of taurine (0.08 mmol) were added to 10 mL of DMSO, stirred magnetically for 12 h, dialyzed and freeze-dried.

The relevant batches were analyzed using HPLC-MS (Agilent 1260 6470 LC/TQ) using a Hilic Amide column (Waters, XBridge BEH Amide 5µm, 4.6x250mm).

### Dose-response against modified β-cyclodextrin

Inhibition assays against hPIV3, Influenza virus, SARS-CoV-2, and RSV were done using a similar protocol based on previously described experiments [17–19].

CDs were serially diluted in DMEM containing for hPIV3 experiments 200 ng/mL TPCK trypsin. Two hundred plaque-forming units (PFU) or infectious units (IFU) of the virus were incubated with the CD dilutions for 1 hour at 37°C. The virus/CD mix was then used to infect host cells—LLCMK2, MDCK-Siat, or Vero E6, depending on the virus plated in an appropriate multi-well plate— in duplicate. After 1 hour incubation at 37°C, the inoculum was removed, and an overlay or maintenance medium was added (DMEM, 0.5% Methylcellulose (MTC) (Sigma), 200 ng/mL TPCK trypsin for hPIV3; DMEM, 2.5% FBS, 0.5% MTC for HSV-2; DMEM, 2.5% FBS, 0.6% Avicel GP3515 (SelectChemie) for SARS-CoV-2; DMEM for Influenza virus and DMEM, 2.5% FBS for RSV-GFP).

Infected cells were incubated at 37°C for 1–5 days depending on the virus. Fixation and staining were performed using crystal violet for hPIV3 and HSV-2; PBS, 4% formaldehyde and water, 20% ethanol, 0.1% crystal violet for SARS-CoV-2; methanol, immunodetection (Ms X Influenza A (1:2000) (Sigma) or anti-influenza B (1:50) (Light Diagnostics) followed by anti-mouse HRP (1:1000) in PBS, 1% BSA, 0.05% Tween-20) and water, 0.1% tablet of 3,3′-Diaminobenzidine, 1.8% H_2_O_2_ for influenza viruses. Plaques for hPIV3, SARS-CoV-2, and HSV-2 or infected cells for influenza viruses and RSV-GFP were quantified manually.

For H7N1, A549 cells (150’000 cells/well) were seeded in 24-well plates. The following day, IAV A/Turkey/Italy/977/1999 (H7N1) modified to express NanoLuc luciferase [70] with a 0.001 MOI were incubated 1 hour with successive dilutions of CD-SLNT/SO3- starting from 100 µg/mL to 0.14µg/mL in Eagle’s Minimum Essential Medium without FBS supplementation. Cells were then infected with the virus:CD-SLNT/SO3- mix for 1 hour. After infection, the inoculum was removed. After 24 hours post-infection, cell layer were lysed in a passive lysis buffer and lysate was used to quantify NanoLuc luciferase activity as previously described [70].

### Virucidal experiment against modified β-cyclodextrin

Virucidal assays against hPIV3, IAV H1N1, SARS-CoV-2, and RSV-GFP were done using a similar protocol based on previously described experiments [17–19].

For hPIV3, 10^5^ plaque-forming units of viruses were incubated with 100 or 300 µg/mL of CDs for 1 or 2 hours at 37°C in DMEM, 200 ng/mL TPCK Trypsin. For Influenza viruses or SARS-CoV-2, 10^5^ ifu or 5 x 10^5^ pfu respectively were incubated with 300 µg/mL of CD-SLNT/SO3- for 1 hour at 37°C in DMEM. The virus/CD mix was serially diluted in DMEM before infection of host cells (LLCMK2, MDCK-Siat, or Vero E6, depending on the virus plated in an appropriate multi-well plate). After 1 hour incubation at 37°C, the inoculum was removed, and an overlay or maintenance medium was added (DMEM, 0.5% Methylcellulose (MTC) (Sigma), 200 ng/mL TPCK trypsin for hPIV3; DMEM, 2.5% FBS, 0.5% MTC for HSV-2; DMEM, 2.5% FBS, 0.6% Avicel GP3515 for SARS-CoV-2; DMEM for Influenza virus and DMEM, 2.5% FBS for RSV-GFP).

Infected cells were incubated at 37°C for 1–5 days depending on the virus. Fixation and staining were performed using crystal violet for hPIV3 and HSV-2; PBS, 4% formaldehyde and water, 20% ethanol, 0.1% crystal violet for SARS-CoV-2; methanol, immunodetection (Ms X Influenza A (1:2000) or anti-influenza B (1:50) followed by anti-mouse HRP (1:1000) in PBS, 1% BSA, 0.05% Tween-20) and water, 0.1% tablet of 3,3′-Diaminobenzidine, 1.8% H_2_O_2_ for influenza viruses. Plaques for hPIV3, SARS-CoV-2, and HSV-2 or infected cells for influenza viruses and RSV-GFP were quantified manually to determine the titer.

### Ex vivo human upper respiratory tract experiments

MucilAir from a pool of donors were purchased from Epithelix and were maintained according to the manufacturer’s protocol using their MucilAir medium. The different viruses used for ex vivo experiments (hPIV3, RSV-GFP, IAV H1N1, or SARS-CoV-2) were produced on MucilAir.

MucilAir were washed with PBS with calcium and magnesium (PBS++) (Gibco) for 45 min at 37°C. Viruses (5x10^4^ to 10^5^ RNA copies) were incubated for 1 hour at 37°C with CD-SLNT/SO3- (300 µg/mL) in MucilAir medium. After incubation, viruses with or without CD were added apically to infect MucilAir for 3 hours at 33°C on the apical side. MucilAir were then washed twice with PBS++ and maintained at 33°C. Every day, an apical wash was performed by adding MucilAir medium at the apical side for 20 min at 33°C. A sample of each apical wash was then used for RT-qPCR.

### Toxicity assays in vitro

LLCMK2 cells were seeded in 96 well plate (10’000 cells/well). CD were serially diluted in DMEM with 200 ng/mL TPCK Trypsin and added to the cells. Cells were incubated at 37°C with the different CDs either for 1 hour before removal and addition of DMEM, 200 ng/mL TPCK Trypsin, or 5 days without removal nor further addition of CDs the subsequent days. After 5 days, cells were washed once with DMEM. A solution of 0.5 mg/mL 3-(4,5-dimethylthiazol-2-yl)-2,5- diphenyltetrazolium bromide (MTT) (Sigma) in DMEM was added for 3 hours at 37°C. Cells were then lysed with DMSO. Absorbance at 570 nm was quantified with a Microplate reader.

### Toxicity assays ex vivo

MucilAir were apically treated daily with 30 µg of CD-SLNT/SO3- and maintained at 33°C. Apical washes were done every day with MucilAir medium for 20 min at 33°C. The basal medium was replaced every day. CyQUANT LDH Cytotoxicity Assay (ThermoFisher) was used according to the manufacturer’s protocol on apical washes. CellTiter 96 Aqueous Cell Proliferation Assay (3-(4,5- dimethylthiazol-2-yl)-5-(3-carboxymethoxyphenyl)-2-(4-sulfophenyl)-2H-tetrazolium (MTS) (Promega) was performed on the apical side of MucilAir according to manufacturer’s protocol.

Resistance against modified β-cyclodextrin For hPIV3, LLCMK2 were seeded on a 6-well plate (300’000 cells / well). HPIV3 clinical #2 was used to infect LLCMK2 in quadruplicates with an MOI of 0.01. After 1 hour at 37°C, the virus inoculum was removed. CD-SLNT/SO3- (50 µg/mL) were added onto two infected wells diluted in DMEM with 200 ng/mL TPCK trypsin. After 5 days post-infection, the supernatant was collected and centrifuged at 2’000 rpm for 5 min. Viruses were titered on LLCMK2 cells. Subsequent passages were performed with the previous virus (MOI 0.01) and CD concentrations were doubled. At passage 10, CD-SLNT/SO3- used was at a concentration of 400 µg/mL. Antiviral activity of CD- SLNT/SO3- against viruses produced at that last passage.

For IAV H1N1, MDCK-Siat were seeded on a 24-well plate (80’000 cells/well). IAV H1N1 was used to infect MDCK-Siat in quadruplicates with an MOI of 0.01. After 1 hour at 37°C, the virus inoculum was removed. CD-SLNT/SO3- (0.2 µg/mL) were added onto two infected wells diluted in DMEM with 1 µg/mL TPCK trypsin. After 2 days post-infection, the supernatant was collected and centrifuged at 2’000 rpm for 5 min. Viruses were titered on MDCK-Siat cells. Subsequent passages were performed with the previous virus (MOI 0.01) and CD concentrations were doubled. At passage 10, CD-SLNT/SO3- used was at a concentration of 230.4 µg/mL. Antiviral activity of CD- SLNT/SO3- against viruses produced at that last passage.

The resistant Influenza A H1N1 viruses were plaque purified by infecting a 6-well plate of MDCK-Siat (300’000 cells/well) in the presence of 300 µg/mL CD-SLNT/SO3- diluted in DMEM, 0.5% MTC, 1 µg/mL TPCK Trypsin. After 2 days post-infection, MDCK-Siat plated on 24 well plates (80’000 cells/well) were infected with a plaque in DMEM, 1 µg/mL TPCK Trypsin. Viruses were collected after 3 to 4 days, tittered, and antiviral activity against CD-SLNT/SO3- was evaluated. Two resistant purified viruses were selected, one per treated passage 10 virus replicates.

### RT-PCR and RT-qPCR

For ex vivo experiments, a sample of each apical wash was lysed with TRK Lysis, and total RNA was extracted with E.Z.N.A total RNA extraction (Omega Bio-Tek) according to the manufacturer’s protocol. RT-qPCR was then performed with TaqPath 1-Step RT-qPCR (ThermoFisher) according to the manufacturer’s protocol with forward/reverse primers and probe specific for each virus (Vi06439670_s1 (ThermoFisher) for hPIV3, 5’- CTCAATTTCCTCACTTCTCCAGTGT-3’/5’- CTTGATTCCTCGGTGTACCTCTGT-3’ and 5’- TCCCATTATGCCTAGGCCAGCAGCA-3’ for RSV-GFP, 5’-GACCRATCCTGTCACCTCTGAC-3’/5’-AGGGCATTYTGGACAAAKCGTCTA-3’ and 5’-TGCAGTCCTCGCTCACTGGGCACG-3’ for IAV H1N1 and 5’ACAGGTACGTTAATAGTTAATAGCGT-3’/5’-ATATTGCAGCAGTACGCACACA-3’ and 5’- ACACTAGCCATCCTTACTGCGCTTCG-3’ for SARS-CoV-2).

For the sequencing of resistant IAV H1N1, the two resistant viruses, the untreated viruses of passage 10 and the original virus were lysed with TRK Lysis, and total RNA was extracted with E.Z.N.A total RNA extraction according to the manufacturer’s protocol. Reverse transcription was performed with SuperScript II (Thermo Fisher) according to the manufacturer’s protocol using random primers. PCR was then performed with Kapa2G (Sigma) according to the manufacturer’s protocol using as primers 5’-GGCAATACTAGTAGTTCTG-3’/5’-CCGTACCATCCATCTACC-3’ and 5’-TGTAAAACGACGGCCAGTCCGAGATATGCATTCGCAATG-3’/5’- CAGGAAACAGCTATGACC TTAAATACATATTCTACACTGTAGAG-3’ for hemagglutinin or 5’- TGTAAAACGACGGCCAGTATGAATCCAAACCAAAAGATAATAAC-3’/5’- CAGGAAACAGCTATGACCCAGGAGCATTCCTCATAGTG-3’ and 5’- TGTAAAACGACGGCCAGTGGTTCTTGCTTTACTGTAATGACC-3’/5’-CAGGAAACAGCTATGACCTTACTTGTCAATGGTAAATGGCAAC-3’ for the neuraminidase of IAV H1N1. Microsynth performed sequencing. The analysis of mutation frequency in GISAID database was done as described previously [71].

### Docking and Molecular dynamics

For SLNT docking, the glycan was created on MOE 2019.0102. PDB 5B2D [39] was prepared on Maestro Schrodinger 2021-2 using the protein preparation wizard generating het states with Epik at a pH of 7.0 ± 2.0 and restrained hydrogens only with OPLS4 force field during minimization. SLNT was prepared using LigPrep by generating possible states with Epik at pH 7.4 while retaining specified chiralities. It was then docked with Glide on the PDB 5B2D using extra precision without post-docking minimization and the 3-sialyl lactose present in the PDB 5B2D as a reference for grid generation using a 20 Å and 56 Å inner and outer cubic boxes respectively. Results were then inspected visually.

For CD-MUS interaction map, PDB 4MZA [42] was prepared on Maetro Schrodinger similarly. CD- MUS were created on MOE and then placed 10 times facing sulfonate groups in front of the protein without touching the protein. Molecular dynamics was then prepared for Desmond simulation by creating an orthorhombic box filled with tip3p water with 0.15 mM NaCl. The molecular dynamics was then run for 500 ns at 300 K and 1.01325 bar. Interactions with sulfonate groups with the protein were extracted and analyzed using Python and Jupyter Notebook. Visualization of the interaction map was done with UCSF ChimeraX 2022-05-29.

For CD-SLNT/SO3- molecular dynamic, the docked SLNT was extracted to MOE. While maintaining this structure intact, the rest of the CD-SLNT/SO3- ligand were created before incorporating the resulting complete ligand into Maestro Schrodinger. PDB 5B2D with CD- SLNT/SO3- having the SLNT groups placed at the sialic acid binding pocket was used as the initial structure for the molecular dynamic. An orthorhombic box was created filled with tip3p water and 0.15 mM NaCl. Desmond was then used for a molecular dynamic of 200 ns at 300 K and 1.01325 bar.

### Antiviral activity of CD-SLNT/SO3- in vivo

Female BALB/c mice were purchased at 7 weeks old from the Centre d’Elevage R. Janvier (Le Genest Saint-Isle, France). Mice were housed in negative-pressure isolators in a containment level 2 facility. Food and water were available ad libitum. At 8 weeks of age, mice were anesthetized with a mixture of ketamine and xylazine (60 mg/kg and 12 mg/kg, respectively per mouse) and infected intranasally with RSV-Luc (10^5^ pfu) as previously performed [11, 72]. CD-SLNT/SO3- were administered as pre-treatment (5 mg/kg/day) intranasally 10 min before infection followed by a daily dose or as post-treatment (10 mg/kg/day) starting 1-day post-infection with a daily dose under anesthesia of 100% oxygen at a rate of 1.5–2 L min−1 mixed with around 4–5% (v/v) isoflurane delivered to the anesthesia chamber (XGI-8, Caliper Life Sciences, Hopkinton, MA, USA). Luminescence measurements were performed at 2 and 4 days post-infection 5 min following instillation of 50 μL D-luciferin (30 mg/ml, Perking Elmer). Living Image software (version 4.0, Caliper Life Sciences) was used to measure the luciferase activity with an exposure time of 1 min. Digital false-color photon emission images of mice were generated and show the average radiance (p/s/cm2/sr). Signals are expressed as total normalized flux (p/s). Mice were euthanized on day 4 post-infection.

### Zebrafish maintenance

Wild type AB adult zebrafish were maintained in the aquatic facility of the KU Leuven at a temperature of 28°C under a 14/10 h light/dark cycle. Fertilized eggs were collected from adults placed in mating cages and were kept in petri dishes containing Danieau’s solution (1.5 mM HEPES, 17.4 mM NaCl, 0.21 mM KCl, 0.12 mM MgSO_4_, 0.18 mM Ca(NO_3_)_2_ and 0.6 μM methylene blue) at 28°C until the start of experiments.

### IAV infection, CD-SLNT/SO3- and baloxavir treatment of zebrafish larvae

Zebrafish larvae at 4.5 days post-fertilization (dpf), i.e. once their swimbladder is inflated, were anesthetized using tricaine (Sigma-Aldrich) and positioned laterally in an agarose mold. Prior to each experiment, the injection needle was calibrated to ensure precise injection volumes. Microinjections were conducted using an M3301R Manual Micromanipulator (WPI) coupled with a FemtoJet 4i pressure microinjector (Eppendorf) [44, 73, 74]. Each larva received a 3 nL injection into the inflated swimbladder of a mixture containing IAV A/Virginia/ATCC3/2009 H1N1 and CD- SLNT/SO3- at a concentration of 100 μg/mL dissolved in absolute DMSO, resulting in ∼20 to 80 of viral RNA copies injected. Post-injection, the larvae were transferred to 6-well plates containing Danieau’s solution and maintained in an incubator set to a 14/10-hour light/dark cycle at 32°C. For baloxavir treatment experiments, baloxavir acid (MedChemExpress) was dissolved in aboslute DMSO and added to the swimming water at a final concentration of 25 μM as described in [44], immediately after IAV H1N1 injection.

### RNA extraction and viral RNA quantification from zebrafish larvae

Each day post-injection, 10 zebrafish larvae were collected into 2 mL tubes containing 1.4 mm ceramic beads (Omni International) and immediately stored at -80°C for subsequent analysis. For RNA extraction, 350 µL of Trizol™ Reagent (Thermo Fisher Scientific) was added to the frozen larvae within the microtubes. The samples were homogenized for 10 seconds at 6’300 rpm using a homogenizer (Bertin Technologies). After homogenization, the lysate was clarified by centrifugation at 100’000 g for 2 minutes, and the supernatant was carefully transferred to a sterile Eppendorf tube. An equal volume of absolute ethanol was mixed with the supernatant, and viral RNA was extracted using the Direct-zol™ RNA Miniprep Kit (Zymo Research), following the manufacturer’s instructions. The purified RNA was stored at -80°C until further quantification by RT-qPCR. Detection of viral RNA was carried out using a one-step RT-qPCR protocol with the iTaq Universal Probes One-Step Kit (Bio-Rad). The primers used were 5’- GACCRATCCTGTCACCTCTGAC-3’ and 5’-AGGGCATTYTGGACAAAKCGTCTA-3’, along with the probe 5’-6-FAM/TGCAGTCCT/ZEN/CGCTCACTGGGCACG/IABkFQ-3’ [75]. Amplification was performed on a QuantStudio™ 5 instrument (Thermo Fisher Scientific) with the following cycling conditions: reverse transcription at 50°C for 10 minutes, initial denaturation at 95°C for 2 minutes, and 40 cycles of 95°C for 15 seconds followed by 60°C for 30 seconds. For absolute quantification, standard curves were generated using 10-fold serial dilutions of a DNA template with a known concentration and the sequence 5’- CTAAAGACAAGACCAATCTTGTCACCTCTGACTAAGGGAATTTTAGGATTTGTGTTCACGCTCACCGTGCCCAGTGAGCGAGGACTGCAGCGTAGACGCTTTGTCCAAAATGCCCTAAATGGGAATGGGGACCC-3’.

## Statistics

In vitro experiments were performed at least in duplicate with at least two independent experiments. Ex vivo experiments were performed with at least two MucilAir in independent experiments. Results are shown as the mean and standard error of the mean (SEM). Half maximal effective concentration (EC_50_) and 50% cytotoxicity concentration (CC_50_) were calculated with GraphPad Prism version 9.1 as described in Mathez and Cagno [68]. ANOVA followed by multiple comparison analysis, unpaired two-tailed t-test, area under the curve followed by additional statistical test, and Mann-Whitney test were used using GraphPad Prism version 9.1. For each figure, the number of independent experiments as well as the statistical test used if applicable are written.

## Supplementary information

The following supporting information is in the joint document: Synthesis of cyclodextrin derivatives, Figure S1 Attachment receptor used by hPIV3, Figure S2 Use of sialic acid and heparan sulfate for hPIV3 infection, Figure S3 Assessing the optimal composition of glycan on β-cyclodextrin, Figure S4 Toxicity of modified β-cyclodextrin, Table S1 Synergy score of combination CD-MUS & CD- SLNT, Figure S5 SLNT, CD-MUS, and CD-SLNT/SO3- interactions with the hemagglutinin-neuraminidase, Figure S6 Virucidal activity of CD-SLNT/SO3-, Figure S7 Virucidal kinetics activity of CD-SLNT/SO3-, Figure S8 Chromatographs of different batches of CD-SLNT/SO3-, Table S2 Influenza A H1N1 mutations after resistance to CD-SLNT/SO3-, Figure S9 In vivo activity of Baloxavir acid, Figure S10 In vivo activity of CD-SLNT/SO3-.

## Data Availability

The data underlying this study are openly available at https://doi.org/10.6084/m9.figshare.27604650.v7. In silico data files are available upon request due to file size limitations.

## Ethics

This study was carried out in accordance with INRAE guidelines in compliance with European animal welfare regulation. The protocols were approved by the Animal Care and Use Committee at “Centre de Recherche de Jouy-en-Josas” (COMETHEA) under relevant institutional authorization (“Ministère de l’éducation nationale, de l’enseignement supérieur et de la recherche”). Authorization number: 2020010910262184 v4 (APAFIS#25172). Zebrafish experiments were approved and performed according to the rules and regulations of the Ethical Committee of KU Leuven (P070/2021), in compliance with the European Union regulations concerning the welfare of laboratory animals as declared in Directive 2010/63/EU.

## Funding and Acknowledgments

This work has been supported by the Swiss National Science Foundation (grant number PZ00P3_193289) to V.C. F.S thanks the very generous funding of the Werner Siemens Foundation. We thank the Cellular Imaging Facility of the University of Lausanne for their help with the acquisition of the glycan array.

## Author Contributions

Conceptualization, G.M, P.J.S, F.S, and V.Cagno; investigation, G.M (in vitro, ex vivo, in silico, synthesis), P.J.S (synthesis, characterization), V.Carlen (in vitro), C.D (in vitro, in vivo), C.P (in vivo), A.H (in vivo), M.G (in vivo), R.LG (in vivo), V.Cagno (in vitro, ex vivo); writing—original draft preparation, G.M, P.J.S, C.D, A.H, J.RP, M.G, R.LG, F.S, and V.Cagno; writing—review and editing, G.M, P.J.S, A.H, J.RP, M.G, R.LG, F.S, and V.Cagno; visualization, G.M, A.H and R.LG; supervision, P.J.S, J. RP, M.G, R.LG, F.S, and V.Cagno; project administration, V.Cagno; funding acquisition, F.S, and V.Cagno. All authors have read and agreed to the published version of the manuscript.

## References

1. Leung Nancy H. L. Transmissibility and transmission of respiratory viruses. Nature Reviews Microbiology. 2021;19(8):528–45.

2. Wang Chia C., Prather Kimberly A., Sznitman Josué, Jimenez Jose L., Lakdawala Seema S., Tufekci Zeynep, Marr Linsey C. Airborne transmission of respiratory viruses. Science. 2021;373(6558):eabd9149.

3. Zhou P., Yang X. L., Wang X. G., Hu B., Zhang L., Zhang W., Si H. R., Zhu Y., Li B., Huang C. L., Chen H. D., Chen J., Luo Y., Guo H., Jiang R. D., Liu M. Q., Chen Y., Shen X. R., Wang X., Zheng X. S., Zhao K., Chen Q. J., Deng F., Liu L. L., Yan B., Zhan F. X., Wang Y. Y., Xiao G. F., Shi Z. L. A pneumonia outbreak associated with a new coronavirus of probable bat origin. Nature. 2020;579(7798):270-3.

4. Uyeki T. M., Milton S., Abdul Hamid C., Reinoso Webb C., Presley S. M., Shetty V., Rollo S. N., Martinez D. L., Rai S., Gonzales E. R., Kniss K. L., Jang Y., Frederick J. C., De La Cruz J. A., Liddell J., Di H., Kirby M. K., Barnes J. R., Davis C. T. Highly Pathogenic Avian Influenza A(H5N1) Virus Infection in a Dairy Farm Worker. N Engl J Med. 2024;390(21):2028–9.

5. Morse J., Coyle J., Mikesell L., Stoddard B., Eckel S., Weinberg M., Kuo J., Riner D., Margulieux K., Stricklen J., Dover M., Kniss K. L., Jang Y., Kirby M. K., Frederick J. C., Lacek K. A., Davis C. T., Uyeki T. M., Lyon-Callo S., Bagdasarian N. Influenza A(H5N1) Virus Infection in Two Dairy Farm Workers in Michigan. N Engl J Med. 2024;391(10):963–4.

6. Mathez G., Cagno V. Viruses Like Sugars: How to Assess Glycan Involvement in Viral Attachment. Microorganisms. 2021;9(6).

7. Conzelmann C., Müller J. A., Perkhofer L., Sparrer K. M., Zelikin A. N., Münch J., Kleger A. Inhaled and systemic heparin as a repurposed direct antiviral drug for prevention and treatment of COVID-19. Clin Med (Lond). 2020;20(6):e218–e21.

8. Frediansyah A. The antiviral activity of iota-, kappa-, and lambda-carrageenan against COVID-19: A critical review. Clin Epidemiol Glob Health. 2021;12:100826.

9. Eccles R., Winther B., Johnston S. L., Robinson P., Trampisch M., Koelsch S. Efficacy and safety of iota-carrageenan nasal spray versus placebo in early treatment of the common cold in adults: the ICICC trial. Respir Res. 2015;16:121.

10. Figueroa J. M., Lombardo M. E., Dogliotti A., Flynn L. P., Giugliano R., Simonelli G., Valentini R., Ramos A., Romano P., Marcote M., Michelini A., Salvado A., Sykora E., Kniz C., Kobelinsky M., Salzberg D. M., Jerusalinsky D., Uchitel O. Efficacy of a Nasal Spray Containing Iota-Carrageenan in the Postexposure Prophylaxis of COVID-19 in Hospital Personnel Dedicated to Patients Care with COVID-19 Disease. Int J Gen Med. 2021;14:6277–86.

11. Cagno Valeria, Andreozzi Patrizia, D’Alicarnasso Marco, Jacob Silva Paulo, Mueller Marie, Galloux Marie, Le Goffic Ronan, Jones Samuel T., Vallino Marta, Hodek Jan, Weber Jan, Sen Soumyo, Janeček Emma-Rose, Bekdemir Ahmet, Sanavio Barbara, Martinelli Chiara, Donalisio Manuela, Rameix Welti Marie-Anne, Eleouet Jean-Francois, Han Yanxiao, Kaiser Laurent, Vukovic Lela, Tapparel Caroline, Král Petr, Krol Silke, Lembo David, Stellacci Francesco. Broad-spectrum non-toxic antiviral nanoparticles with a virucidal inhibition mechanism. Nature Materials. 2018;17(2):195–203.

12. Cagno V., Tseligka E. D., Jones S. T., Tapparel C. Heparan Sulfate Proteoglycans and Viral Attachment: True Receptors or Adaptation Bias? Viruses. 2019;11(7).

13. Guillon P., Dirr L., El-Deeb I. M., Winger M., Bailly B., Haselhorst T., Dyason J. C., von Itzstein M. Structure-guided discovery of potent and dual-acting human parainfluenza virus haemagglutinin-neuraminidase inhibitors. Nat Commun. 2014;5:5268.

14. Pascolutti M., Dirr L., Guillon P., Van Den Bergh A., Ve T., Thomson R. J., von Itzstein M. Structural Insights into Human Parainfluenza Virus 3 Hemagglutinin-Neuraminidase Using Unsaturated 3- N-Substituted Sialic Acids as Probes. ACS Chem Biol. 2018;13(6):1544–50.

15. Ali Eunüs S., Rajapaksha Harinda, Carr Jillian M., Petrovsky Nikolai. Norovirus drug candidates that inhibit viral capsid attachment to human histo-blood group antigens. Antiviral Research. 2016;133:14–22.

16. Heida Rick, Bhide Yoshita C., Gasbarri Matteo, Kocabiyik Özgün, Stellacci Francesco, Huckriede Anke L. W., Hinrichs Wouter L. J., Frijlink Henderik W. Advances in the development of entry inhibitors for sialic-acid-targeting viruses. Drug Discovery Today. 2021;26(1):122–37.

17. Jones S. T., Cagno V., Janecek M., Ortiz D., Gasilova N., Piret J., Gasbarri M., Constant D. A., Han Y., Vukovic L., Kral P., Kaiser L., Huang S., Constant S., Kirkegaard K., Boivin G., Stellacci F., Tapparel C. Modified cyclodextrins as broad-spectrum antivirals. Sci Adv. 2020;6(5):eaax9318.

18. Kocabiyik Ozgun, Cagno Valeria, Silva Paulo Jacob, Zhu Yong, Sedano Laura, Bhide Yoshita, Mettier Joelle, Medaglia Chiara, Da Costa Bruno, Constant Samuel, Huang Song, Kaiser Laurent, Hinrichs Wouter L. J., Huckriede Anke, Le Goffic Ronan, Tapparel Caroline, Stellacci Francesco. Non-Toxic Virucidal Macromolecules Show High Efficacy Against Influenza Virus Ex Vivo and In Vivo. Advanced Science. 2021;8(3):2001012.

19. Cagno V., Gasbarri M., Medaglia C., Gomes D., Clement S., Stellacci F., Tapparel C. Sulfonated Nanomaterials with Broad-Spectrum Antiviral Activity Extending beyond Heparan Sulfate-Dependent Viruses. Antimicrob Agents Chemother. 2020;64(12).

20. Gasbarri Matteo, V’kovski Philip, Torriani Giulia, Thiel Volker, Stellacci Francesco, Tapparel Caroline, Cagno Valeria. SARS-CoV-2 Inhibition by Sulfonated Compounds. Microorganisms [Internet]. 2020; 8(12).

21. Chandrasekaran A., Srinivasan A., Raman R., Viswanathan K., Raguram S., Tumpey T. M., Sasisekharan V., Sasisekharan R. Glycan topology determines human adaptation of avian H5N1 virus hemagglutinin. Nat Biotechnol. 2008;26(1):107–13.

22. Bose Santanu, Banerjee Amiya K. Role of Heparan Sulfate in Human Parainfluenza Virus Type 3 Infection. Virology. 2002;298(1):73–83.

23. Amonsen Mary, Smith David F., Cummings Richard D., Air Gillian M. Human Parainfluenza Viruses hPIV1 and hPIV3 Bind Oligosaccharides with α2-3-Linked Sialic Acids That Are Distinct from Those Bound by H5 Avian Influenza Virus Hemagglutinin. Journal of Virology. 2007;81(15):8341–5.

24. Tappert M. M., Smith D. F., Air G. M. Fixation of oligosaccharides to a surface may increase the susceptibility to human parainfluenza virus 1, 2, or 3 hemagglutinin-neuraminidase. J Virol. 2011;85(23):12146–59.

25. Suzuki T., Portner A., Scroggs R. A., Uchikawa M., Koyama N., Matsuo K., Suzuki Y., Takimoto T. Receptor specificities of human respiroviruses. J Virol. 2001;75(10):4604–13.

26. Fukushima Keijo, Takahashi Tadanobu, Ito Seigo, Takaguchi Masahiro, Takano Maiko, Kurebayashi Yuuki, Oishi Kenta, Minami Akira, Kato Tatsuya, Park Enoch Y., Nishimura Hidekazu, Takimoto Toru, Suzuki Takashi. Terminal sialic acid linkages determine different cell infectivities of human parainfluenza virus type 1 and type 3. Virology. 2014;464–465:424-31.

27. Moscona A., Porotto M., Palmer S., Tai C., Aschenbrenner L., Triana-Baltzer G., Li Q. X., Wurtman D., Niewiesk S., Fang F. A recombinant sialidase fusion protein effectively inhibits human parainfluenza viral infection in vitro and in vivo. J Infect Dis. 2010;202(2):234–41.

28. Zwygart A. C., Medaglia C., Zhu Y., Bart Tarbet E., Jonna W., Fage C., Le Roy D., Roger T., Clement S., Constant S., Huang S., Stellacci F., Silva P. J., Tapparel C. Development of Broad-Spectrum beta-Cyclodextrins-Based Nanomaterials Against Influenza Viruses. J Med Virol. 2024;96(12):e70101.

29. Ianevski A., Giri A. K., Aittokallio T. SynergyFinder 3.0: an interactive analysis and consensus interpretation of multi-drug synergies across multiple samples. Nucleic Acids Res. 2022;50(W1):W739–43.

30. Yadav B., Wennerberg K., Aittokallio T., Tang J. Searching for Drug Synergy in Complex Dose-Response Landscapes Using an Interaction Potency Model. Comput Struct Biotechnol J. 2015;13:504–13.

31. Tang J., Wennerberg K., Aittokallio T. What is synergy? The Saariselkä agreement revisited. Front Pharmacol. 2015;6:181.

32. Bliss C. I. THE TOXICITY OF POISONS APPLIED JOINTLY1. Annals of Applied Biology. 1939;26(3):585–615.

33. Loewe S., Muischnek H. Über Kombinationswirkungen. Naunyn-Schmiedebergs Archiv für experimentelle Pathologie und Pharmakologie. 1926;114(5):313–26.

34. Loewe S. Die quantitativen Probleme der Pharmakologie. Ergebnisse der Physiologie. 1928;27(1):47–187.

35. Vlot Anna H. C., Aniceto Natália, Menden Michael P., Ulrich-Merzenich Gudrun, Bender Andreas. Applying synergy metrics to combination screening data: agreements, disagreements and pitfalls. Drug Discovery Today. 2019;24(12):2286–98.

36. Porotto M., Fornabaio M., Kellogg G. E., Moscona A. A second receptor binding site on human parainfluenza virus type 3 hemagglutinin-neuraminidase contributes to activation of the fusion mechanism. J Virol. 2007;81(7):3216–28.

37. Yuan P., Thompson T. B., Wurzburg B. A., Paterson R. G., Lamb R. A., Jardetzky T. S. Structural studies of the parainfluenza virus 5 hemagglutinin-neuraminidase tetramer in complex with its receptor, sialyllactose. Structure. 2005;13(5):803–15.

38. Crennell S., Takimoto T., Portner A., Taylor G. Crystal structure of the multifunctional paramyxovirus hemagglutinin-neuraminidase. Nat Struct Biol. 2000;7(11):1068–74.

39. Kubota M., Takeuchi K., Watanabe S., Ohno S., Matsuoka R., Kohda D., Nakakita S. I., Hiramatsu H., Suzuki Y., Nakayama T., Terada T., Shimizu K., Shimizu N., Shiroishi M., Yanagi Y., Hashiguchi T. Trisaccharide containing alpha2,3-linked sialic acid is a receptor for mumps virus. Proc Natl Acad Sci U S A. 2016;113(41):11579–84.

40. Porotto M., Palmer S. G., Palermo L. M., Moscona A. Mechanism of fusion triggering by human parainfluenza virus type III: communication between viral glycoproteins during entry. J Biol Chem. 2012;287(1):778–93.

41. Marcink T. C., Yariv E., Rybkina K., Más V., Bovier F. T., des Georges A., Greninger A. L., Alabi C. A., Porotto M., Ben-Tal N., Moscona A. Hijacking the Fusion Complex of Human Parainfluenza Virus as an Antiviral Strategy. mBio. 2020;11(1).

42. Xu R., Palmer S. G., Porotto M., Palermo L. M., Niewiesk S., Wilson I. A., Moscona A. Interaction between the hemagglutinin-neuraminidase and fusion glycoproteins of human parainfluenza virus type III regulates viral growth in vivo. mBio. 2013;4(5):e00803–13.

43. Gabor K. A., Goody M. F., Mowel W. K., Breitbach M. E., Gratacap R. L., Witten P. E., Kim C. H. Influenza A virus infection in zebrafish recapitulates mammalian infection and sensitivity to anti-influenza drug treatment. Dis Model Mech. 2014;7(11):1227–37.

44. Van Dycke J., Cuvry A., Knickmann J., Ny A., Rakers S., Taube S., de Witte P., Neyts J., Rocha-Pereira J. Infection of zebrafish larvae with human norovirus and evaluation of the in vivo efficacy of small-molecule inhibitors. Nat Protoc. 2021;16(4):1830–49.

45. Simon A. K., Hollander G. A., McMichael A. Evolution of the immune system in humans from infancy to old age. Proc Biol Sci. 2015;282(1821):20143085.

46. Parker Edward P. K., Desai Shalini, Marti Melanie, Nohynek Hanna, Kaslow David C., Kochhar Sonali, O’Brien Katherine L., Hombach Joachim, Wilder-Smith Annelies. Response to additional COVID-19 vaccine doses in people who are immunocompromised: a rapid review. The Lancet Global Health. 2022;10(3):e326–e8.

47. Zwygart Arnaud Charles-Antoine, Medaglia Chiara, Zhu Yong, Tarbet E Bart, Jonna Westover, Fage Clément, Le Roy Didier, Roger Thierry, Constant Samuel, Huang Song, Stellacci Francesco, Silva Paulo Jacob, Tapparel Caroline. Development of Broad-spectrum β-cyclodextrins-Based Nanomaterials Against Influenza Viruses. bioRxiv. 2024.

48. Marcink T. C., Zipursky G., Cheng W., Stearns K., Stenglein S., Golub K., Cohen F., Bovier F., Pfalmer D., Greninger A. L., Porotto M., des Georges A., Moscona A. Subnanometer structure of an enveloped virus fusion complex on viral surface reveals new entry mechanisms. Sci Adv. 2023;9(6):eade2727.

49. DuBois R. M., Aguilar-Yanez J. M., Mendoza-Ochoa G. I., Oropeza-Almazan Y., Schultz-Cherry S., Alvarez M. M., White S. W., Russell C. J. The receptor-binding domain of influenza virus hemagglutinin produced in Escherichia coli folds into its native, immunogenic structure. J Virol. 2011;85(2):865–72.

50. Zhang W., Shi Y., Qi J., Gao F., Li Q., Fan Z., Yan J., Gao G. F. Molecular basis of the receptor binding specificity switch of the hemagglutinins from both the 1918 and 2009 pandemic influenza A viruses by a D225G substitution. J Virol. 2013;87(10):5949–58.

51. Xu R., McBride R., Nycholat C. M., Paulson J. C., Wilson I. A. Structural characterization of the hemagglutinin receptor specificity from the 2009 H1N1 influenza pandemic. J Virol. 2012;86(2):982–90.

52. Zhu Y., Gasbarri M., Zebret S., Pawar S., Mathez G., Diderich J., Valencia-Camargo A. D., Russenberger D., Wang H., Silva P. H. J., Dela Cruz J. B., Wei L., Cagno V., Munz C., Speck R. F., Desmecht D., Stellacci F. Benzene with Alkyl Chains Is a Universal Scaffold for Multivalent Virucidal Antivirals. ACS Cent Sci. 2024;10(5):1012–21.

53. Alymova I. V., Portner A., Mishin V. P., McCullers J. A., Freiden P., Taylor G. L. Receptor-binding specificity of the human parainfluenza virus type 1 hemagglutinin-neuraminidase glycoprotein. Glycobiology. 2012;22(2):174–80.

54. Klimyte E. M., Smith S. E., Oreste P., Lembo D., Dutch R. E. Inhibition of Human Metapneumovirus Binding to Heparan Sulfate Blocks Infection in Human Lung Cells and Airway Tissues. J Virol. 2016;90(20):9237–50.

55. Chang A., Masante C., Buchholz U. J., Dutch R. E. Human metapneumovirus (HMPV) binding and infection are mediated by interactions between the HMPV fusion protein and heparan sulfate. J Virol. 2012;86(6):3230–43.

56. Milewska A., Zarebski M., Nowak P., Stozek K., Potempa J., Pyrc K. Human coronavirus NL63 utilizes heparan sulfate proteoglycans for attachment to target cells. J Virol. 2014;88(22):13221–30.

57. Lang J., Yang N., Deng J., Liu K., Yang P., Zhang G., Jiang C. Inhibition of SARS pseudovirus cell entry by lactoferrin binding to heparan sulfate proteoglycans. PLoS ONE. 2011;6(8):e23710.

58. Shi Y., Zhang W., Wang F., Qi J., Wu Y., Song H., Gao F., Bi Y., Zhang Y., Fan Z., Qin C., Sun H., Liu J., Haywood J., Liu W., Gong W., Wang D., Shu Y., Wang Y., Yan J., Gao G. F. Structures and receptor binding of hemagglutinins from human-infecting H7N9 influenza viruses. Science. 2013;342(6155):243-7.

59. Ramos I., Krammer F., Hai R., Aguilera D., Bernal-Rubio D., Steel J., Garcia-Sastre A., Fernandez-Sesma A. H7N9 influenza viruses interact preferentially with alpha2,3-linked sialic acids and bind weakly to alpha2,6-linked sialic acids. J Gen Virol. 2013;94(Pt11):2417–23.

60. Lee N., Khalenkov A. M., Lugovtsev V. Y., Ireland D. D., Samsonova A. P., Bovin N. V., Donnelly R. P., Ilyushina N. A. The use of plant lectins to regulate H1N1 influenza A virus receptor binding activity. PLoS ONE. 2018;13(4):e0195525.

61. Liu Y., Childs R. A., Matrosovich T., Wharton S., Palma A. S., Chai W., Daniels R., Gregory V., Uhlendorff J., Kiso M., Klenk H. D., Hay A., Feizi T., Matrosovich M. Altered receptor specificity and cell tropism of D222G hemagglutinin mutants isolated from fatal cases of pandemic A(H1N1) 2009 influenza virus. J Virol. 2010;84(22):12069–74.

62. Moss R. B., Hansen C., Sanders R. L., Hawley S., Li T., Steigbigel R. T. A phase II study of DAS181, a novel host directed antiviral for the treatment of influenza infection. J Infect Dis. 2012;206(12):1844–51.

63. Mathez Gregory, Pillonel Trestan, Bertelli Claire, Cagno Valeria. Alpha and Omicron SARS-CoV-2 Adaptation in an Upper Respiratory Tract Model. Viruses. 2023;15(1):13.

64. Varricchio Carmine, Mathez Gregory, Pillonel Trestan, Bertelli Claire, Kaiser Laurent, Tapparel Caroline, Brancale Andrea, Cagno Valeria. Geneticin shows selective antiviral activity against SARS-CoV-2 by interfering with programmed −1 ribosomal frameshifting. Antiviral Research. 2022;208:105452.

65. Essaidi-Laziosi M., Brito F., Benaoudia S., Royston L., Cagno V., Fernandes-Rocha M., Piuz I., Zdobnov E., Huang S., Constant S., Boldi M. O., Kaiser L., Tapparel C. Propagation of respiratory viruses in human airway epithelia reveals persistent virus-specific signatures. J Allergy Clin Immunol. 2018;141(6):2074–84.

66. Essaidi-Laziosi M., Alvarez C., Puhach O., Sattonnet-Roche P., Torriani G., Tapparel C., Kaiser L., Eckerle I. Sequential infections with rhinovirus and influenza modulate the replicative capacity of SARS-CoV-2 in the upper respiratory tract. Emerg Microbes Infect. 2022;11(1):412–23.

67. Heida R., Akkerman R., Jacob Silva P. H., Lakerveld A. J., Ortiz D., Bigogno C., Gasbarri M., van Kasteren P. B., Stellacci F., Frijlink H. W., Huckriede A. L. W., Hinrichs W. L. J. Development of an inhalable antiviral powder formulation against respiratory syncytial virus. J Control Release. 2023;357:264–73.

68. Mathez G., Cagno V. Clinical severe acute respiratory syndrome coronavirus 2 isolation and antiviral testing. Antivir Chem Chemother. 2021;29:20402066211061063.

69. Hallak L. K., Collins P. L., Knudson W., Peeples M. E. Iduronic acid-containing glycosaminoglycans on target cells are required for efficient respiratory syncytial virus infection. Virology. 2000;271(2):264–75.

70. Mettier J., Marc D., Sedano L., Da Costa B., Chevalier C., Le Goffic R. Study of the host specificity of PB1-F2-associated virulence. Virulence. 2021;12(1):1647–60.

71. Franzi E., Mathez G., Dinant S., Deloizy C., Kaiser L., Tapparel C., Le Goffic R., Cagno V. Non-Steroidal Estrogens Inhibit Influenza Virus by Interacting with Hemagglutinin and Preventing Viral Fusion. Int J Mol Sci. 2023;24(20).

72. Sake S. M., Zhang X., Rajak M. K., Urbanek-Quaing M., Carpentier A., Gunesch A. P., Grethe C., Matthaei A., Ruckert J., Galloux M., Larcher T., Le Goffic R., Hontonnou F., Chatterjee A. K., Johnson K., Morwood K., Rox K., Elgaher W. A. M., Huang J., Wetzke M., Hansen G., Fischer N., Eleouet J. F., Rameix-Welti M. A., Hirsch A. K. H., Herold E., Empting M., Lauber C., Schulz T. F., Krey T., Haid S., Pietschmann T. Drug repurposing screen identifies lonafarnib as respiratory syncytial virus fusion protein inhibitor. Nat Commun. 2024;15(1):1173.

73. Van Dycke J., Dai W., Stylianidou Z., Li J., Cuvry A., Roux E., Li B., Rymenants J., Bervoets L., de Witte P., Liu H., Neyts J., Rocha-Pereira J. A Novel Class of Norovirus Inhibitors Targeting the Viral Protease with Potent Antiviral Activity In Vitro and In Vivo. Viruses. 2021;13(9).

74. Van Dycke J., Ny A., Conceicao-Neto N., Maes J., Hosmillo M., Cuvry A., Goodfellow I., Nogueira T. C., Verbeken E., Matthijnssens J., de Witte P., Neyts J., Rocha-Pereira J. A robust human norovirus replication model in zebrafish larvae. PLoS Pathog.2019;15(9):e1008009.

75. Prevention Centers for Disease Control and. CDC protocol of realtime PCR for influenza A(H1N1)2009. 2009.

