## Supporting Information for "A pan-respiratory virus attachment inhibitor with high potency in human airway models and in vivo"

**Supporting Information Text**

**Synthesis of cyclodextrin derivatives.**

Cyclodextrin Core β-cyclodextrin(-S-C11-COOH)7 [1, 2]: All reagents were dried before use under ~2.5x10-2mbar at room temperature for 48 hours; dry solvents were purchased and used as received. Potassium tert-butoxide (tBUOK, 82.47 mmol: 8.7g, Sigma) and 12-mercaptododecanoic acid (35.28 mmol: 8.2 g, Sigma) were dissolved in 100 mL of dry DMF under an argon atmosphere and the mixture was stirred at room temperature using a mechanical stirrer, 300rpm. Heptakis-(6-deoxy-6-iodo)-β-Cyclodextrin (4.2 mmol: 8 g, Arachem) was added to the mixture after one hour, the mixture was placed in an oil bath at 70°C and the reaction proceeded overnight (12 hours). The crude, was precipitated into 1L of diethyl ether (Et2O) and vacuum filtered with a fritted disk funnel (POR 3). The solid on the filter was collected (25 g), dissolved in 190 mL of distilled water. Then 380 mL of ACN was added to the aqueous solution creating an off-white homogeneous suspension. This mixture was vacuum filtered (POR4) – the crude was allowed to percolate through the filter by gravity: once a flow of solvent was observed by gravity alone, the vacuum assisted filtration was turned on and the filtration went smoothly. ~12 grams of an off-white solid was collected and dried. It was again dissolved in 200mL of pure water and precipitated into 0.1M HCl, to obtain the protonated species as a white powder, collected via centrifugation in 45mL Falcon tubes, followed by decantation. 1H NMR (400 MHz, TFA-d) δ 5.37 (d, J = 3.6 Hz, 1H, glucose H-1), 4.35 (t, J = 9.0 Hz, 1H, glucose H-3), 4.28 (t, J = 7.4 Hz, 1H, glucose H-5), 4.10 (dd, J = 9.9, 3.4 Hz, 1H, glucose H-2), 3.88 (t, J = 9.3 Hz, 1H, glucose H-4), 3.48 (d, J = 12.8 Hz, 1H, glucose H-6a), 3.24 (dd, J = 13.5, 7.6 Hz, 1H, glucose H-6b), 3.00–2.84 (m, 2H, S-CH2-CH2), 2.64 (t, J = 7.6 Hz, 2H, CH2-COOH), 1.86 (q, J = 7.4 Hz, 4H, CH2-CH2-COOH, S-CH2-CH2), 1.53 (m, 14H, S-C-C-CH2-CH2-CH2-CH2-CH2-CH2-CH2-C-C-COO). HRMS (nanochip-ESI/LTQ-Orbitrap) m/z: [M–3H]3− Calcd for C126H221O42S73− 877.5133; Found 877.1148.

Activated β-cyclodextrin(-S-CD-NHS)7 [1, 2]: β-CD-(-S-C11-COOH)7 (0.377 mmol: 10 g) was dissolved in 100 mL DMSO under magnetic stirring until a transparent homogeneous solution formed. Then, N-hydroxysuccinimide (5.317 mmol: 1.03 g, Sigma Aldrich) was added followed by the addition of 1-Ethyl-3-(3-dimethylaminopropyl)carbodiimide (9.53 mmol: 1.85 g Sigma Aldrich), and 4-Dimethylaminopyridine (1.22 mmol: 150 mg Sigma Aldrich) and stirred for 12 hours. The crude was washed with cold acidic water (500µL of 1 M HCl into 2L of MilliQ Water kept at 4°C) and centrifuged at 5500 rpm at 4°C for 5 minutes inside 45mL falcon tubes. Generally, four falcon tubes were filled with 30mL of the acidic water. Then, 10mL of the crude was added into the 30mL of acidic water into per tube, forming a white precipitate. The dispersion was centrifuged at 5500 rpm, the supernatant discarded, another 30mL of the acidic water added, followed by the crude, repeating the procedure until the pellet became too large for the falcon tube. The procedure was started using a new falcon tube to complete all the 100mL of crude. Once the solid was collected, additional washes with acidic washes were performed, aiming at 5 washes per pellet. The pellets were redispersed using a sonicator bath in alternation with vigorous shaking and vortexing. As the number of washes progressed, the time of centrifugation was extended to 10 or 15min as needed to sediment all the visible solid material. Then, Acetonitrile (ACS grade, Sigma Aldrich) was used to wash the white crude. Sonication and vigorous agitation with the aid of a vortex was used to completely redisperse the crude in Acetonitrile and centrifuge it down at 5500 rpm and 4°C. This was repeated until a cloudy white supernatant that could not be further sedimented under these conditions was formed. This translucent (opaque) off-white supernatant was discarded and the wash was finished using Diethyl Ether (ACS grade, Fisher Scientific). The crude was dispersed in Et2O using vigorous agitation, vortexing and sonication when needed. The Et2O washed were done 5 times to completely remove the acetonitrile The final pellets were dried inside a desiccator under high vacuum for 12 hours and collected as a dry powder (~12 grams).

Characterization of relevant batches using HPLC-MS. HPLC-MS (Agilent 1260 6470 LC/TQ) using a Hilic Amide column (Waters, XBridge BEH Amide 5µm, 4.6x250mm) following the gradient below was performed for each sample. A is MilliQ water with 10mM of ammonium formate, pH 3 and B is mass grade Acetonitrile.

| Time | A (%) | B (%) | Flow (mL/min) |
|---|---|---|---|
| 0 | 15 | 85 | 1 |
| 30 | 30 | 70 | 1 |
| 65 | 40 | 60 | 1 |
| 70 | 40 | 60 | 1 |
| 85 | 45 | 55 | 1 |
| 95 | 50 | 50 | 1 |
| 105 | 60 | 40 | 1 |
| 110 | 60 | 40 | 1 |

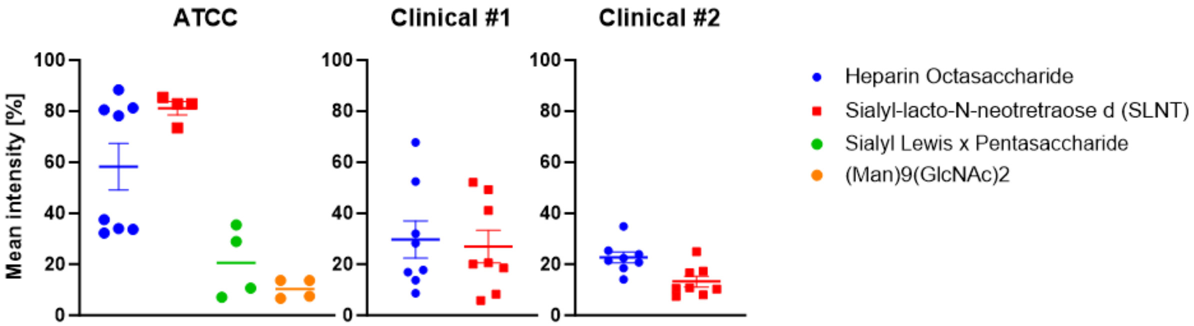


**Figure S1 Attachment receptor used by hPIV3.** Laboratory strain (ATCC) and two clinical isolates of hPIV3 were incubated overnight at 4°C on a glycan array that displayed 100 different glycans. Intensities of fluorescence are expressed relative to the fluorescence of the positive control and background. Data represent mean ± SEM (Standard Error of the Mean) of two independent experiments.

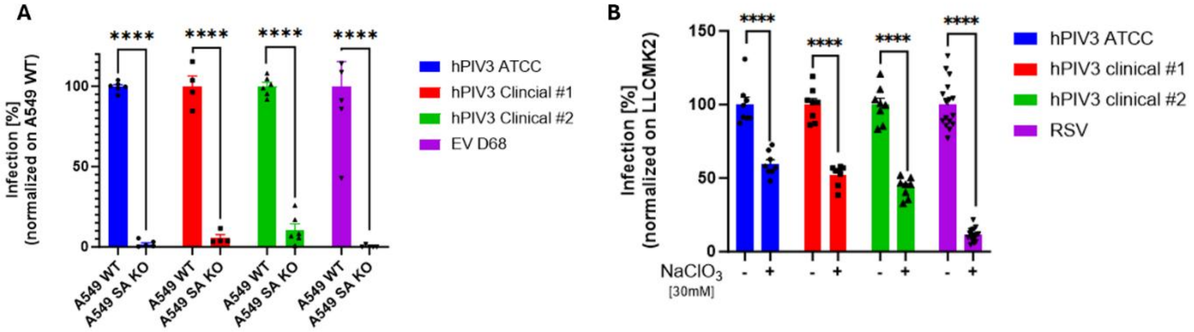


**Figure S2 Use of sialic acid and heparan sulfate for hPIV3 infection.** (A) A549 with sialic acid transporter knockout (A549 SA KO) were infected with hPIV3 and EV68. After immunostaining, the level of infectivity was evaluated by tetramethyl benzidine. Data represent mean ± SEM of three independent experiments. Two-tailed t-tests were performed to compare WT and SA KO data for each virus. (B) LLCMK2 cells were passaged in the presence or not of 30 mM sodium chlorate ($NaClO_3$). They were then infected with hPIV3 or RSV-GFP. Infectivity was assessed by plaque assay for hPIV3 and number of infectious units for RSV-GFP. Data represent mean ± SEM of two independent experiments. Two-tailed t-tests were performed to compare untreated and treated for each virus. * P< 0.0332, ** P<0.0021, *** P<0.0002, **** P<0.0001

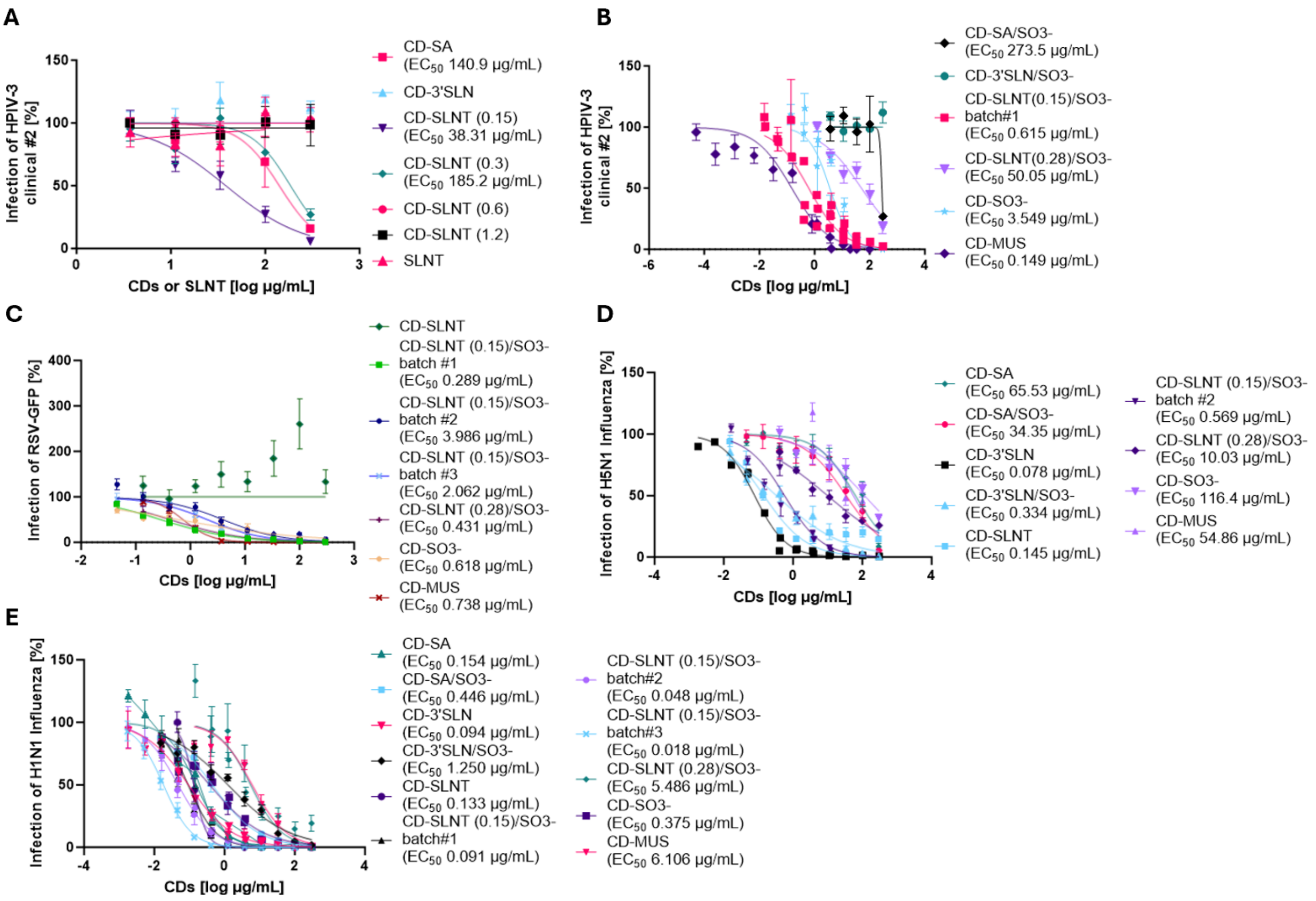


**Figure S3 Assessing the optimal composition of glycan on β-cyclodextrin.** (A) Antiviral activity of CD-SLNT with different equivalences (0.15, 0.3, 0.6, 1.2 i.e amount of SLNT grafted onto the modified β-cyclodextrin), CD-3'SLN, CD-SA, or SLNT alone against Human Parainfluenza virus 3 (hPIV-3) clinical #2 were performed. (B) Sulfated versions of these cyclodextrins (CD-SA/SO3-, CD-3'SLN/SO3-, CD-SLNT/SO3- with two different equivalences of SLNT), CD-MUS or CD-SO3- (cyclodextrin synthesized similarly to CD-SLNT/SO3- without SLNT) were tested against HPIV-3 clinical #2.(C) CD-SLNT and its respective sulfated version (CD-SLNT/SO3-), CD-MUS and CD-SO3- were tested against RSV-GFP. (D-E) CD-SA, CD-3'SLN, CD-SLNT and their respective sulfated versions (CD-SA/SO3-, CD-3'SLN/SO3-, CD-SLNT/SO3- respectively), CD-MUS and CD-SO3- were tested against Influenza A H5N1 (D) or H1N1 (E) viruses. The pre-incubation was done for 1 hour at 37°C before infection on LCCMK2 cells for HPIV-3 clinical #2 (A-B), or RSV-GFP (C) and on MDCK-Siat for Influenza A H5N1 (D) or H1N1 (E) viruses. Data represents mean ± SEM of two (A – CD-SA, CD-3'SLN, CD-SLNT (0.6, 1.2), SLNT; B – CD-SA/SO3-, CD-3'SLN/SO3-; C – CD-SLNT, CD-SLNT(0.28)/SO3-, CD-SO3-, CD-MUS; D – CD-SA, CD-SA/SO3-, CD-3'SLN/SO3-, CD-SLNT(0.28)/SO3-, CD-MUS, CD-SO3-; E – CD-3'SLN/SO3-, CD-SLNT, CD-MUS), three (A –CD-SLNT (0.15, 0.3); C- CD-SLNT (0.15)/SO3- batch#2, CD-SLNT (0.15)/SO3- batch#3; D – CD-SLNT (0.15)/SO3- batch#2, CD-3'SLN, E - CD-SA, CD-3'SLN, CD-SO3-), four (B – CD-SLNT(0.28)/SO3-; C – CD-SLNT (0.15)/SO3- batch#1; D – CD-SLNT, E – CD-SA/SO3-, CD-SLNT (0.15)/SO3- batch#3, CD-SLNT(0.28)/SO3-), five (B – CD-SO3-, CD-SLNT (0.15)/SO3- batch#1; E - CD-SLNT (0.15)/SO3- batch#1, #2) or six (B – CD-MUS) independent experiments. Nonlinear regression with variable Hill Slope and constraint bottom and top (0 and 100 respectively) were performed to compute $EC_{50}$.

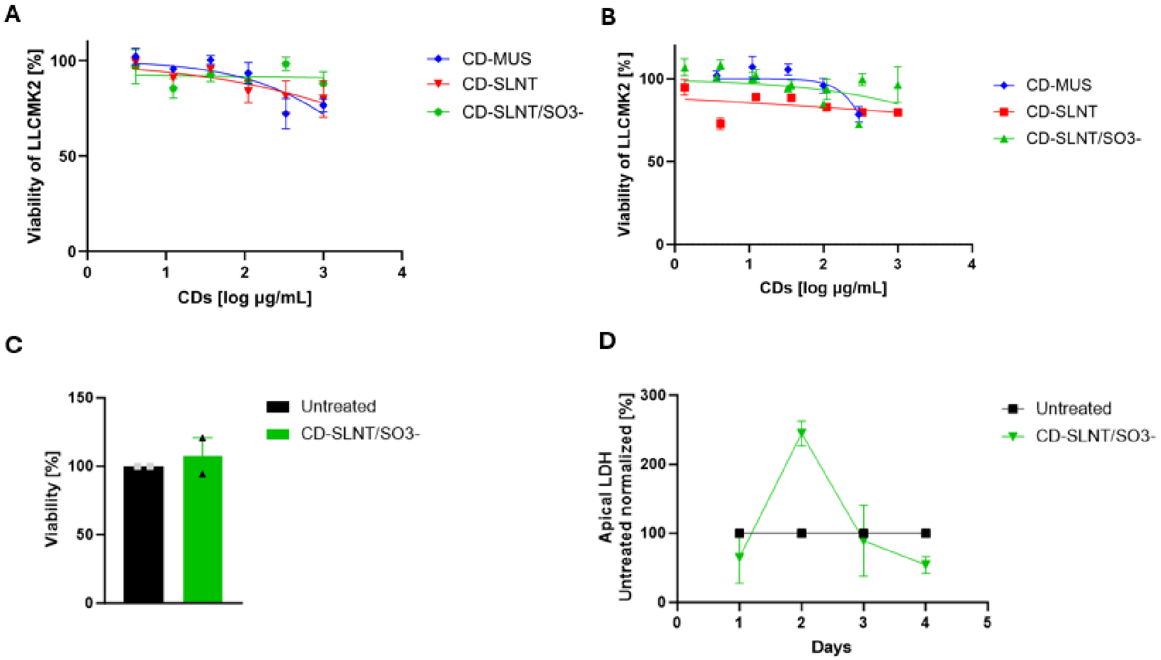


**Figure S4 Toxicity of modified β-cyclodextrin.** Toxicity of CD-MUS, CD-SLNT, and CD-SLNT/SO3- was assessed in vitro and ex vivo. CDs were incubated for 1h (A) or 5 days (B) on LLCMK2. Cell viability was quantified five days post-exposure by MTT. Data represents mean ± SEM of two (A, B – CD-SLNT) or five independent experiments (B – CD-MUS, CD-SLNT/SO3-). Nonlinear regression with variable Hill slope and constraints for the bottom and top (0 and 100 respectively) were performed to compute $CC_{50}$. (C, D) CDs were administered apically in human respiratory airways daily (n = 2 for each condition). (C) Viability was assessed by MTS 4 days after the first exposure. Data represents mean ± SEM. A two-tailed t-test was performed. (D) LDH was quantified on the daily apical of the human respiratory airway. Data represents mean ± SEM. The statistical analysis was performed by calculating the area under the curve followed by a two-tailed t-test.

| Virus | HSA | Loewe | Bliss | ZIP |
|---|---|---|---|---|
| hPIV3 ATCC | -33.662 ± 20.41 | -51.465 ± 20.41 | -34.894 ± 20.41 | -32.199 ± 20.41 |
| hPIV3 clinical #1 | -6.223 ± 6.58 | -9.525 ± 6.58 | -14.206 ± 6.58 | -10.41 ± 6.58 |
| hPIV3 clinical #2 | -11.876 ± 10.93 | -28.393 ± 10.93 | -14.054 ± 10.93 | -12.491 ± 10.93 |

**Table S1 Synergy score of combination CD-MUS & CD-SLNT**

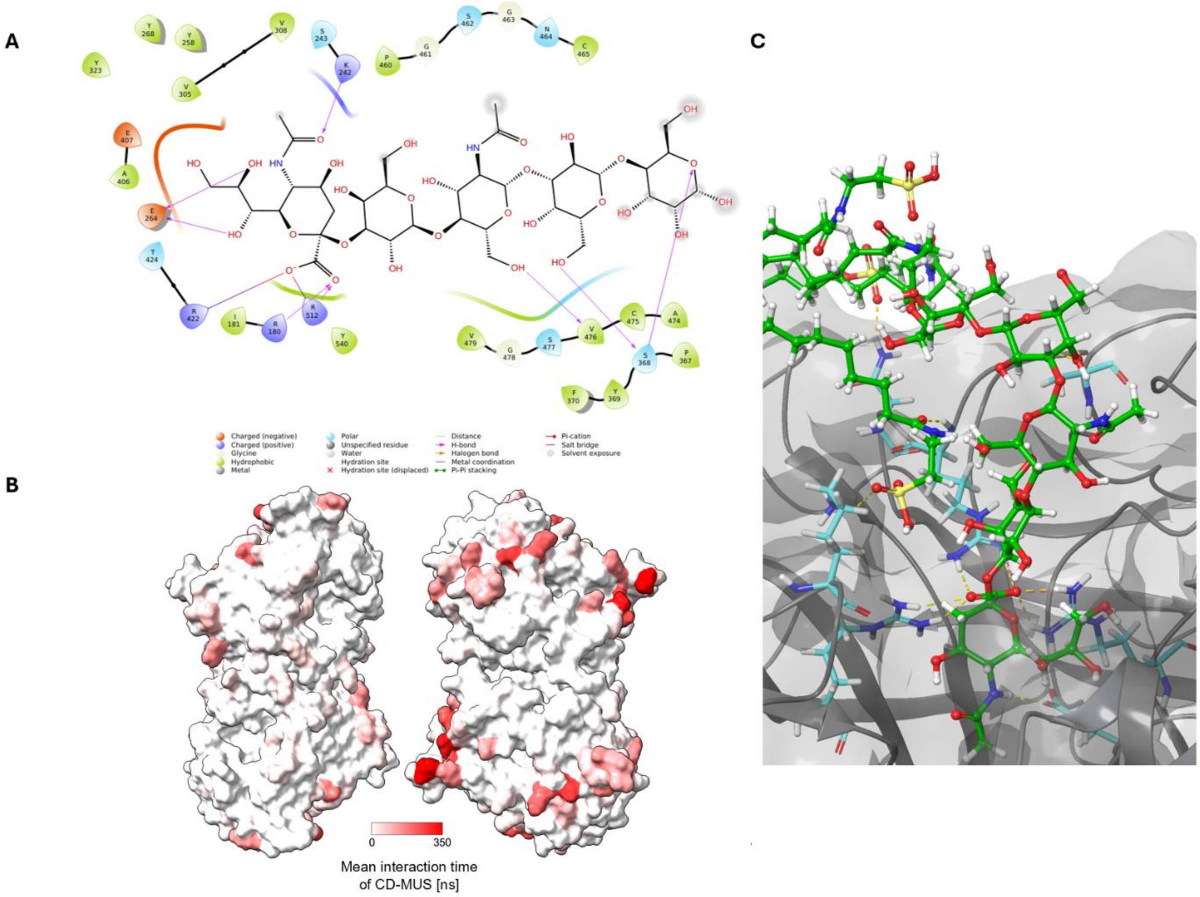

**Figure S5 SLNT, CD-MUS, and CD-SLNT/SO3- interactions with the hemagglutinin-neuraminidase.** (A) SLNT was docked on the hemagglutinin-neuraminidase (PDB : 5B2D). An interaction map is presented. (B) Interactions of CD-MUS with the hemagglutinin-neuraminidase (PDB : 4MZA) were simulated by molecular dynamics. Results are aggregates of two molecular dynamics simulations of 500 ns with 10 CD-MUS. Each CD-MUS were located at a different position around the protein. The nanomaterial was facing the active ligands to the protein. The first 200 ns of the simulation was excluded of the analysis to avoid potential bias of the initial position. (C) Representation of the interaction of CD-SLNT/SO3- (green) with residues of the hemagglutinin-neuraminidase (cyan) (PDB : 5B2D) during a molecular dynamic of 200 ns.

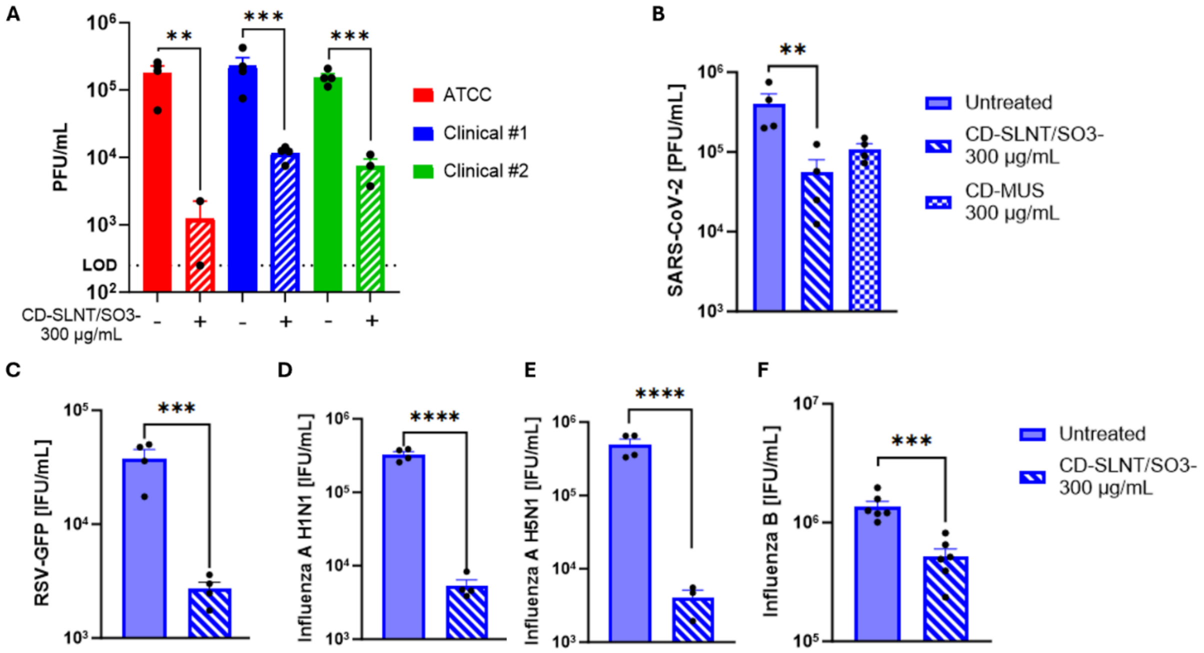


**Figure S6 Virucidal activity of CD-SLNT/SO3-.** HPIV3 (A), SARS-CoV-2 (B), RSV (C), Influenza A H1N1 (D), Influenza A H5N1 (E) or Influenza B (F) were incubated with 300 µg/mL of CD-SLNT/SO3- for 1 hour at 37°C. Viral titers were then quantified. Data represents mean ± SEM of two (A,B,C,D,E) or three (F) independent experiments. Two-tailed t-tests were performed to compare untreated and treated conditions. * P< 0.0332, ** P<0.0021, *** P<0.0002, **** P<0.0001

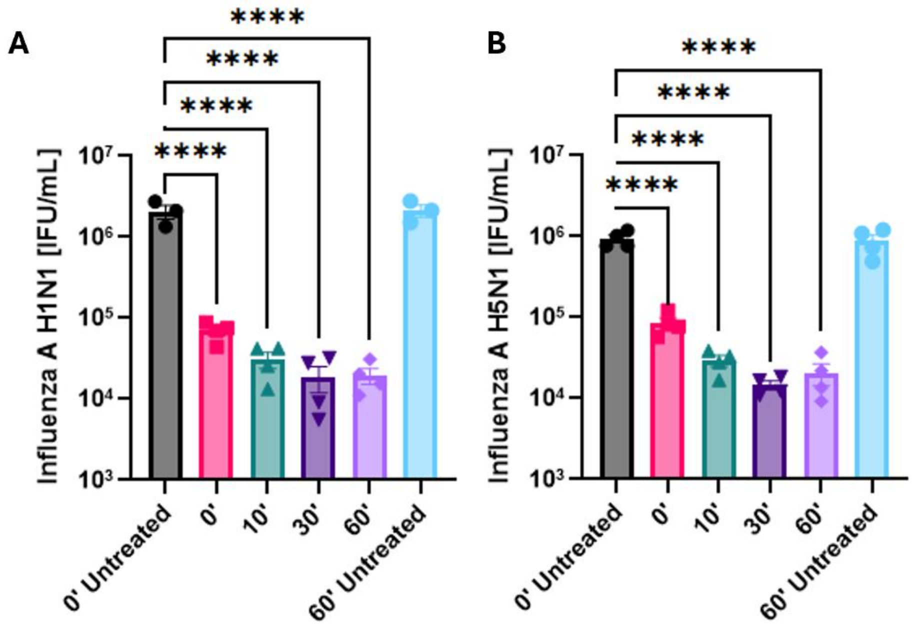


**Figure S7 Virucidal kinetics activity of CD-SLNT/SO3-.** IAV H1N1 (A) and IAV H5N1 (B) were incubated with 300 µg/mL of CD-SLNT/SO3- for 0, 10, 30, or 60 minutes at 37°C. Viral titers were then quantified. Data represents mean ± SEM of two independent experiments. One-way ANOVA with Dunnett's multiple comparisons compared to 0' Untreated was performed to compare the different conditions. **** P<0.0001

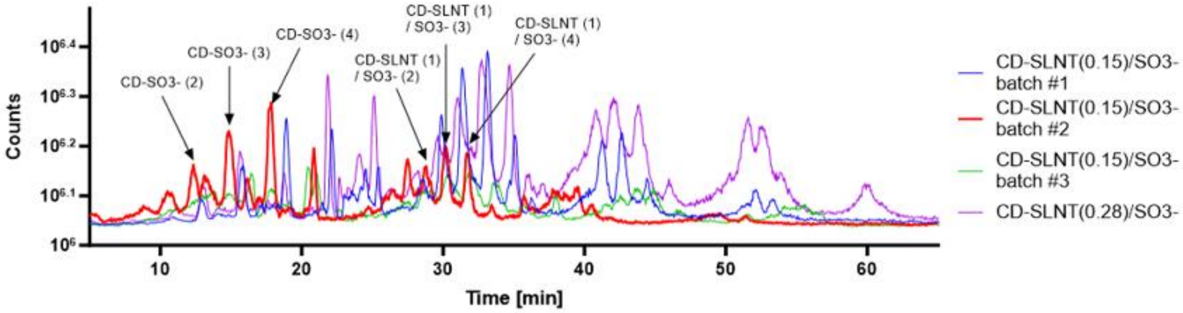


**Figure S8 Chromatographs of different batches of CD-SLNT/SO3-.** High-performance liquid chromatography coupled with mass spectrometry was performed with the different produced batch of CD-SLNT/SO3-. Arrows identified pics for which mass spectrometry was confident with a type of species that can be synthesized. The analysis showed that batches that mix SLNT and Taurine generate both species with SO3- alone that elute before 20 minutes with this method, and species with SLNT and differing number of taurine groups. Centered around 30 min are the species with 1 SLNT that we hypothesize to be the most potent, between 40-45 minutes species with 2 SLNT and different amounts of SO3- and between 50-55 minutes species with 3 SLNT and different amounts of SO3-.

| Gene | Mutation | Frequency in GISAID |
|---|---|---|
| HA | **K154E[#1]** | 0.008% |
| | **G155E[#2]** | 0.453% |
| | R205K[#1] | 2.436% |
| | N377N/T[#1] | 0.042% |
| | Q452Q/P[#1, #2] | 0% |
| | K458K/Q[#1, #2] | 0.011% |
| | G/V479V[#1, #2] | 99.93%[†] |
| | T483T/N[#2] | 0.037% |
| | D485D/Y[#2] | 0.031% |
| NA | Q/E45Q*[#1, #2] | - |
| | K/Q51Q*[#1, #2] | - |
| | I54N[#2] | - |
| | V80G/V[#2] | - |
| | P/A86A[#1, #2] | - |
| | R/S95S*[#1, #2] | - |
| | V407I/V[#1] | - |

**Table S2 Influenza A H1N1 mutations after resistance to CD-SLNT/SO3-.** Bold HA mutations are closed to the sialic acid binding pocket of the protein. #1, #2 Mutations present in the resistant plaque-purified Influenza A H1N1 viruses #1 or #2 respectively. * Mutations present in one replicate of untreated virus † V479 is the natural amino acid of Influenza A H1N1 hemagglutinin.

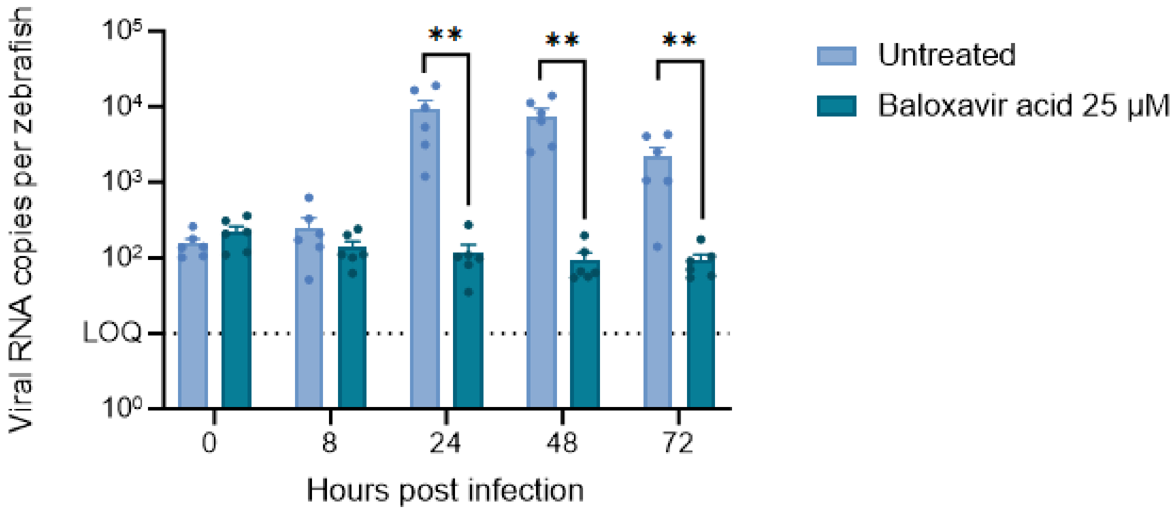


**Figure S9 In vivo activity of Baloxavir acid.** Zebrafish larvae were infected with IAV H1N1. Baloxavir acid was added to the swimming water at a concentration of 25mM. At 8 hpi and each day, 10 larvae are lysed for viral RNA quantification by RT-qPCR. Data represents mean ± SEM of six independent experiments. Two-tailed Mann-Whitney tests were performed to compare untreated and treated groups. ** P<0.0021

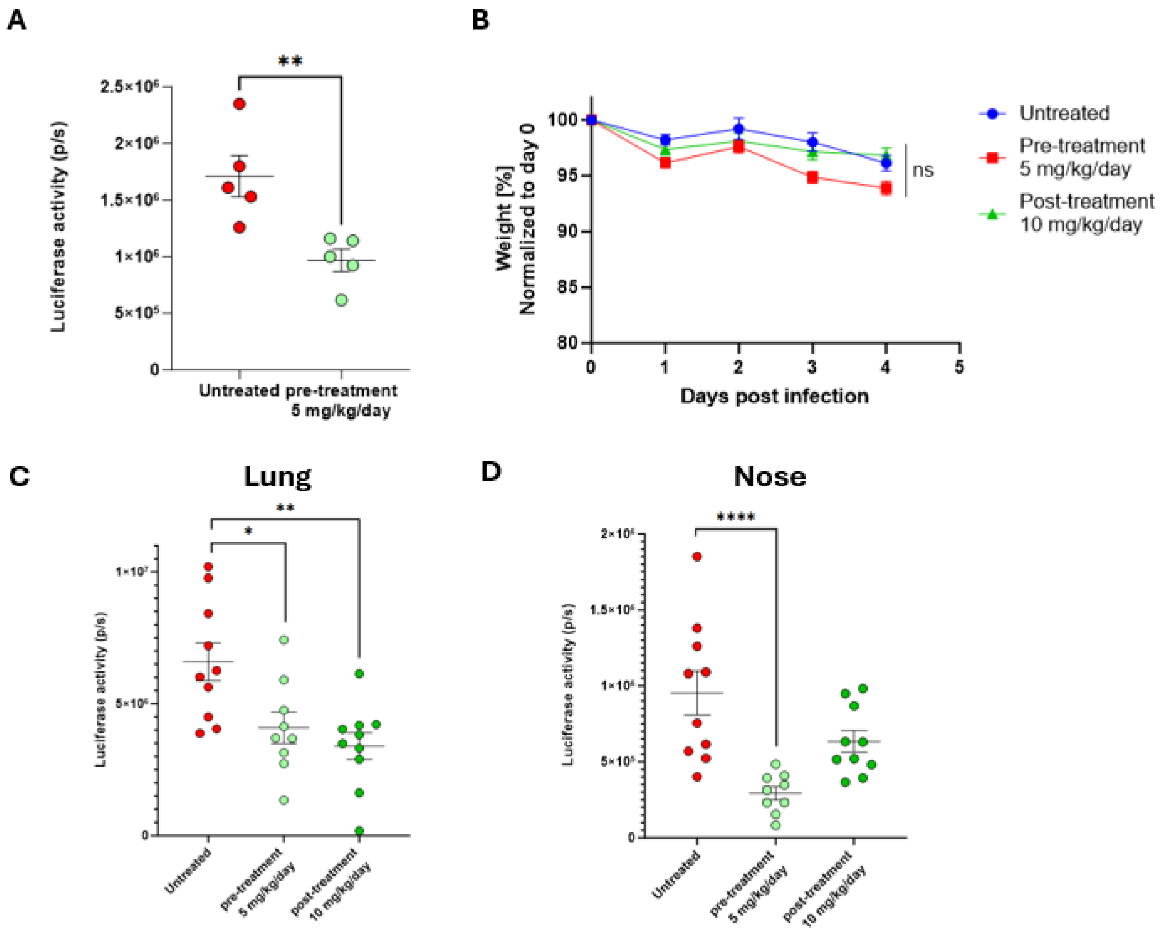


**Figure S10 In vivo activity of CD-SLNT/SO3-.** Mice (n = 5 (A), 10 (B, C, D) per group) were infected with RSV-Luc. Mice were either treated with 5 mg/kg of CD-SLNT/SO3- 30 minutes before infection (pre-treatment) with a daily dose until 4 days post-infection. (A) RSV replication was assessed by luciferase activity on day 4 for the initial experiment. During a subsequent experiment (B, C, D), infected mice were pre-treated similarly as initially or with 10 mg/kg of CD-SLNT/SO3- starting 1-day post-infection (post-treatment) until 4 days post-infection. Mice weights were measured every day (B). RSV replication was assessed by luciferase activity on day 4 in the lungs (C) and nose (D). Data represents mean ± SEM of one experiment. Two-tailed Mann-Whitney tests were performed to compare untreated and each condition. * $P < 0.0332$, ** $P<0.0021$, *** $P<0.0002$, **** $P<0.0001$, non significant (ns)

**SI References**


1. Heida, R., R. Akkerman, P. H. Jacob Silva, A. J. Lakerveld, D. Ortiz, C. Bigogno, M. Gasbarri, P. B. van Kasteren, F. Stellacci, H. W. Frijlink, A. L. W. Huckriede, and W. L. J. Hinrichs. "Development of an Inhalable Antiviral Powder Formulation against Respiratory Syncytial Virus." J Control Release 357 (2023): 264-73.
2. Zwygart, A. C., C. Medaglia, Y. Zhu, E. Bart Tarbet, W. Jonna, C. Fage, D. Le Roy, T. Roger, S. Clement, S. Constant, S. Huang, F. Stellacci, P. J. Silva, and C. Tapparel. "Development of Broad-Spectrum Beta-Cyclodextrins-Based Nanomaterials against Influenza Viruses." J Med Virol 96, no. 12 (2024): e70101.
